# Effects of phylogeny on coexistence in model communities

**DOI:** 10.1101/2020.09.04.283507

**Authors:** Carlos A. Serván, José A. Capitán, Zachary R. Miller, Stefano Allesina

## Abstract

A species’ traits influence the way in which it interacts with the environment. Thus, we expect traits to play a role in determining whether a given set of species coexists. Traits are, in turn, the outcome of an eco-evolutionary process summarized by a phylogenetic tree. Therefore, the phylogenetic tree associated with a set of species should encode information about the assembly properties of the community. Many studies have highlighted the potentially complex ways in which phylogenetic information is translated into species’ ecological properties. However, much less emphasis has been placed on developing expectations for community properties under a particular hypothesis.

In this work, we couple a simple model of trait evolution on a phylogenetic tree with local community dynamics governed by Lotka-Volterra equations. This allows us to derive properties of the community of coexisting species as a function of the number of traits, tree topology and the size of the species pool. Our results highlight how phylogenies and traits, in concert, affect the coexistence of a set of species.

In this way, our work provides new baseline expectations for the ways in which phylogenetic information is reflected in the structure of and coexistence within local communities.

## Introduction

Gause’s pioneering work [15] provided the first clear empirical evidence for the principle of competitive exclusion, which states that two species competing for a unique resource cannot coexist. In the context of niche theory, this principle resonates in the concept of limiting similarity: In a community shaped only by biotic interactions, species with similar niches are less likely to coexist [26]. Making stronger assumptions, one can draw a direct link between evolutionary relatedness among the members of an ecological community and their co-occurrence patterns. In particular, if one is willing to assume that species’ traits are well-described by phylogeny, and that similarity in traits maps into strength of competition between species, one can connect the phylogenetic structure of an ecological community with coexistence [37]. While this hypothesis has found mixed support [11], it has served as one of the cornerstones of the budding field of community phylogenetics [35, 38]. In recent years, several tools have been developed to test whether a given mechanism of community assembly (e.g., competitive exclusion or environmental filtering) has acted on a community, by analyzing the signal it is expected to leave in the community’s phylogenetic structure [14]. However, some authors have noted that phylogenetic relatedness might affect community patterns in a variety of ways, obscuring a link between phylogenetic and co-occurrence patterns [11, 28].

Here we take a step back and analyze a model in which we incorporate an explicit link between phylogenetic relatedness and ecological interactions. In particular, we connect phylogeny to species’ traits, and then similarity in traits to the strength of interaction between any two species [4, 29]. Given a phylogenetic tree representing the evolutionary history of a regional pool of *n* species, we assume that species interactions are determined by a set of *ℓ ≥ n* traits, which have evolved independently on the tree via Brownian motion [18]. Species are assumed to have a baseline competitive effect on each other, which is then modified according their trait covariance. In this way, species that are more closely related tend to interact, on average, more strongly with each other than with distantly-related species. As we will show, the variance of the distribution of interaction strengths is controlled by the number of traits *ℓ*.

Clearly, species’ intrinsic growth rates could also reflect their evolutionary history (e.g., closely related species with similar traits might find similar environments to be harsh or hospitable). To clearly separate the effect of phylogeny on interspecific interactions from its effect on growth rates, we therefore assume that all species have the same intrinsic growth rate. [5]. This assumption severs any connection between phylogeny and environmental filtering.

Having established our model for trait evolution and the link between trait values and species interactions, we analyze the case in which all species in the pool are present at arbitrary initial conditions, and dynamics follow the Generalized Lotka-Volterra model. Contrary to previous simulation-based studies [14, 23] we develop an analytical framework to characterize the resulting community of coexisting species, as a function of both the number of traits, *ℓ*, and the tree structure. In particular, we show that when the number of traits is large enough relative to the number of species in the pool, coexistence of all species is guaranteed by the tree-induced interaction structure. Furthermore, the abundance distribution of the community reflects the structure of the tree. On the other hand, while *ℓ = n* is a well-known *necessary* condition for coexistence [24, 41], we find that full coexistence is almost never achieved in this case (see also [13]). Yet, even when coexistence of all *n* species is precluded, one typically observes coexisting communities of moderate size, as expected if interactions were purely random [10, 33]. Differently from the purely random case, here we find that the probability that a particular species survives is determined by its position in the tree.

Our model shows that phylogenetic relatedness, modulated by the number of traits controlling species interactions, affects multiple aspects of the local community. The explicit incorporation of community dynamics allows us to move from pairwise comparisons to global aspects of community structure. Furthermore, we advance the growing body of literature on random interaction models [3, 6, 10, 33] by analyzing a case in which the correlations between interaction strengths are controlled by phylogenetic relatedness.

## Model

Consider a regional pool 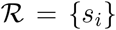 of *n* species indexed by 1 ≤ *i* ≤ *n*, and assume that a species’ identity is defined by its *ℓ ≥ n* trait values. For a given trait 1 ≤ *j ≤ ℓ*, collect the values of *j* for all members of the pool in the trait vector 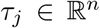. We sample each *τ_j_* independently from a multivariate normal distribution 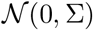. This choice implies that: (a) the values for distinct traits of a given species are independent, and thus we are not considering trade-offs between traits; (b) the processes leading to the correlation structure ∑ are statistically equivalent for distinct traits; (c) lastly, if ∑_*ii*_ = *σ* for all *i*, then the distribution of trait values *within* a species is independent of species identity. For an example of an evolutionary process consistent with the assumptions above, consider the case in which 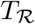 is the phylogenetic tree for the species in the regional pool, and each trait *j* starts at an ancestral mean value of 0, and evolves independently on the tree via Brownian motion. Then the value of trait *j* at the *n* tips, *τ_j_*, follows the normal distribution 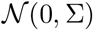 with ∑ induced by the tree structure, and called the variance-covariance matrix of 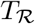 [18]. In ∑, the covariance between two species is given by the shared branch length on 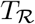 [9]. As such, whenever 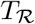 is ultrametric, then ∑_*ii*_ = 1 for all *i*. Unless otherwise specified, here ∑ is always assumed to originate from an ultrametric, rooted phylogenetic tree (see Figures 1 and 2).

**Figure 1:**
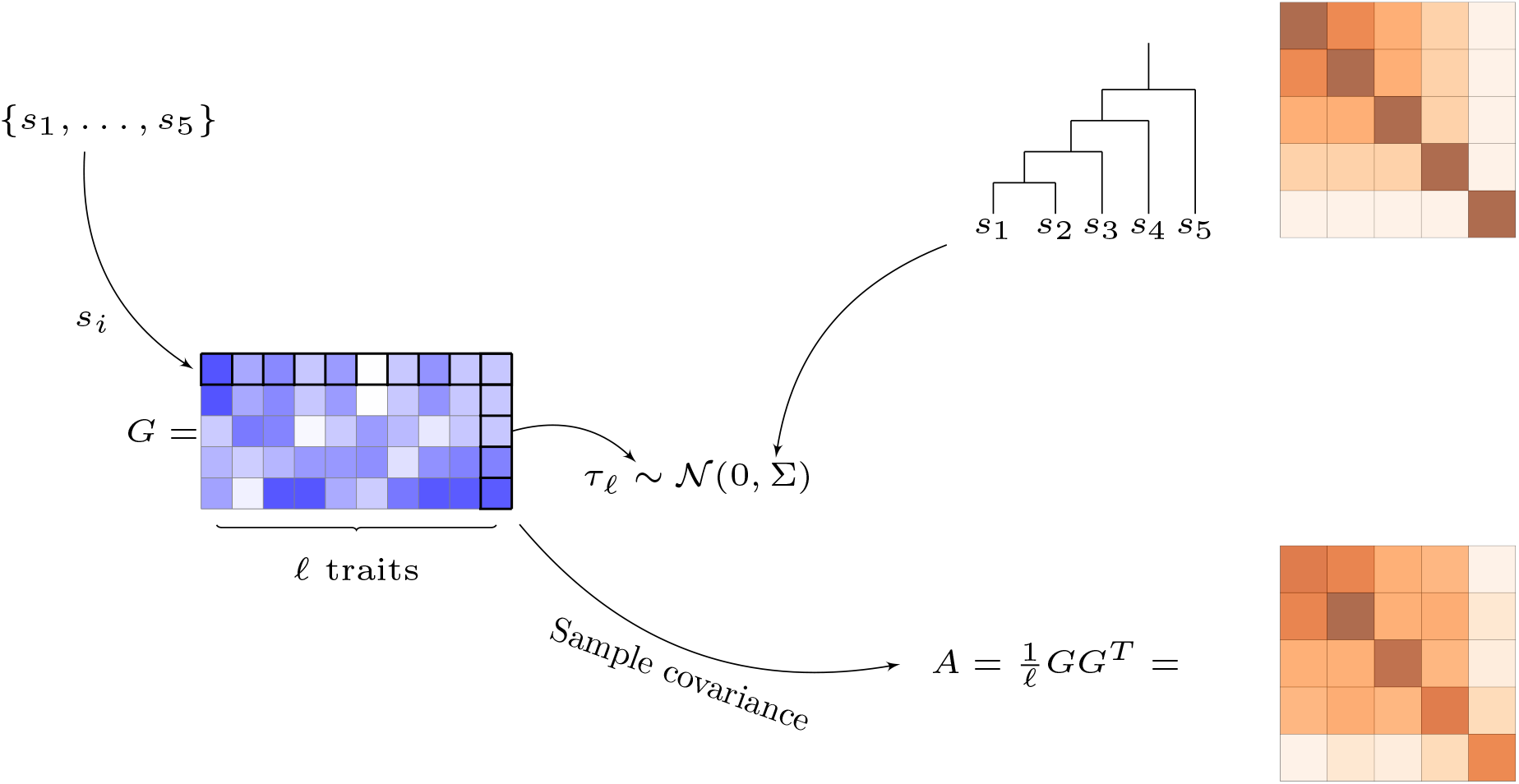
Construction of the regional pool ℛ and interaction matrix *A*. Each species in the pool 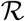 is assigned *ℓ* trait values. The vector containing the values for trait 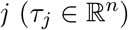 of all members of the pool is sampled independently from 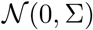. This is equivalent to a neutral model of trait evolution for each *j* on a phylogenetic tree 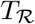. The model relates the structure of 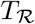 to the interactions between the species in the pool: the matrix ∑ measures the shared evolutionary history between any two species *s_i_* and *s_j_* on 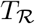 (in our example ∑_12_ > ∑_13_ > … > ∑_15_). In turn, the number of traits *ℓ* and ∑ determine the interactions between species, stored in the matrix *A*.

**Figure 2:**
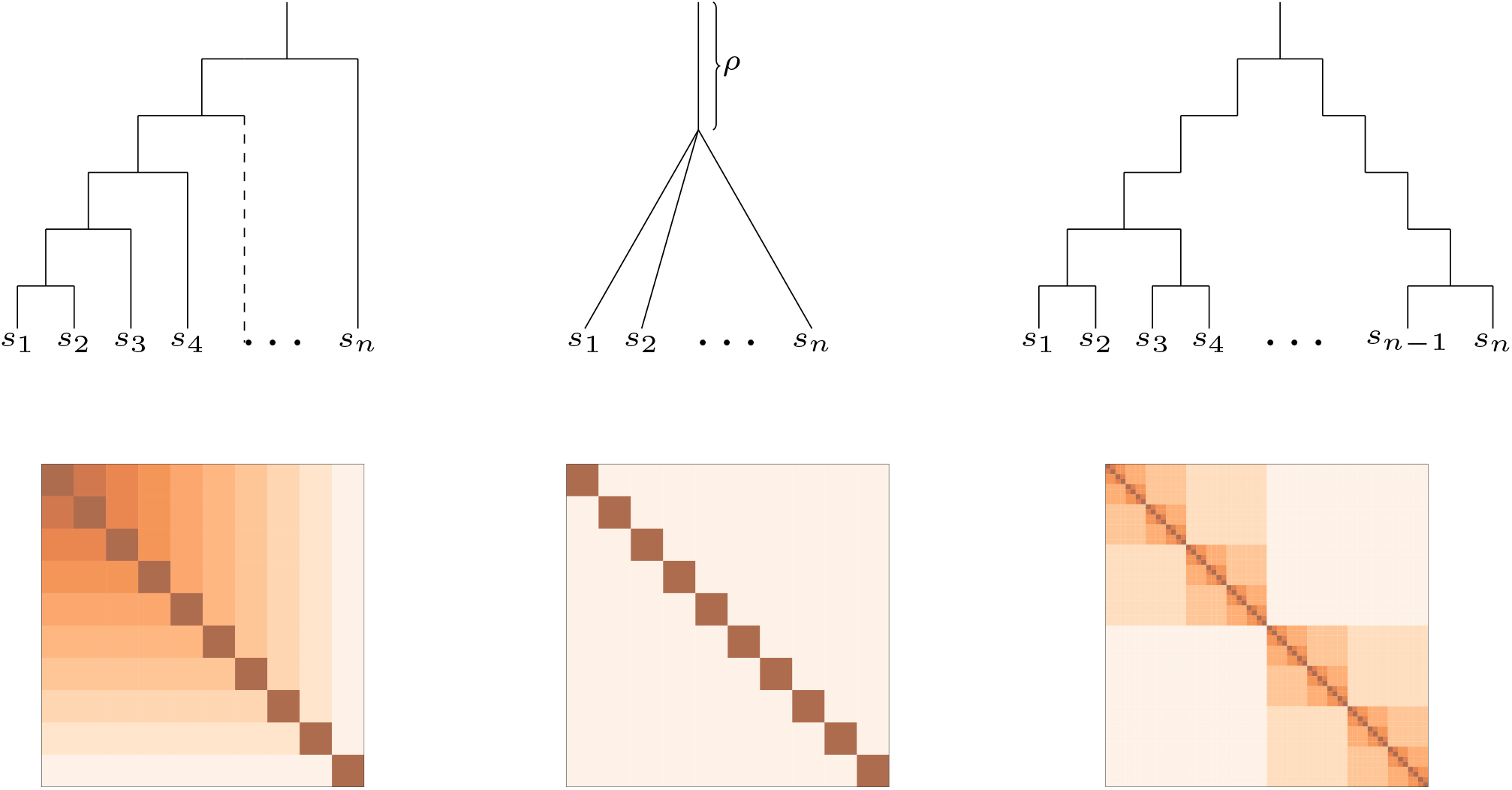
Examples of ultrametric rooted phylogenies and its induced covariance matrices. The perfectly unbalanced tree (left) has *n* − 1 branching times 0 < *t*_1_ < … < *t*_*n*−1_ for a pool of *n* species, where each new branching happens to the “left” and creates a new pair of species. We call the times between branching events, *t_i_ − t*_*i*−1_, *inter-branching times*. The star tree (middle) displays a unique branching event which generates all the *n* species. For the perfectly balanced tree (right) we have *n* branching *times* at each of which all the tips present up to that point generate two new species. Proceeding recursively, *n* branching times generate 2^*n*^ species and we have *n* + 1 distinct inter-branching times. The covariance matrix associated with each tree is constructed as follows: For any *s_i_* take *γ_i_* to be the path “backwards” in time to the ancestral species at the root of the tree, then for any two *s_i_, s_j_*, let *t*(*i,j*) be the time at with *γ_i_* and *γ_j_* merge, i.e., the coalescence time between *s_i_* and *s_j_* [36]. Then, ∑_*ij*_ = 1 − *t*(*i,j*). In particular, ∑_*ij*_ is the total time for which the evolutionary processes for *s_i_* and *s_j_* are completely linked. For example, in the star tree ∑_*ij*_ = *ρ* for any *i = j* and ∑_*ii*_ = 1, given that each tree is ultrametric.

In this setting, each realization of the *ℓ* traits defines a species pool 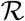. For a given pool, we imagine that the following experiment is performed [33]: all the species from the pool are introduced in the local habitat *at the same time* and at *arbitrary initial densities*. Population dynamics, as determined by the species’ interactions and growth rates, will lead the community to an *asymptotic* state in which some of the species are extinct, while others coexist. Our aim is to characterize the resulting community of coexisting species in terms of the parameters *ℓ, n* and ∑.

To this end, we consider dynamics governed by the Generalized Lotka-Volterra (GLV) model. Species are assumed to differ only in their interactions, so that the growth of each species in isolation is the same:

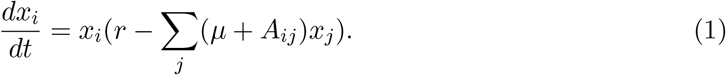

Here, *x_i_* is the density of species *i* and *r* is the common intrinsic growth rate. The interaction coefficients are modeled as deviations from a “mean-field” competition value *μ* > 0. These deviations are controlled by trait similarity between species. More precisely, the deviations are modeled as the sample covariance matrix resulting from the trait sampling process, so that competition between two species is strengthened if their trait vectors are positively correlated and weakened otherwise:

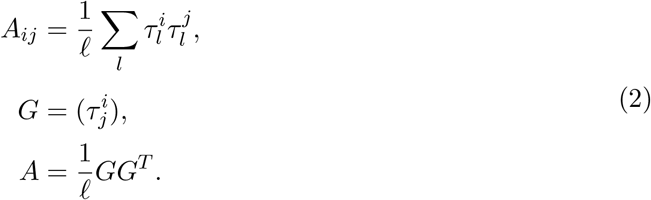

In the supplementary information (section S1), we show how this model arises by assuming a separation of time scales for consumer-resource models in which consumers share the same attack and death rates, but differ in their preferences for resources.

Under these assumptions, *A* is a symmetric and stable matrix, and a member of the *Wishart ensemble* [30, 40]:

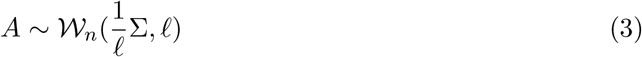

The Wishart distribution describes the probability with which a given *sample covariance matrix* is observed when sampling from a multivariate normal distribution. Given its many applications in statistics and other fields, the Wishart distribution has been studied extensively, allowing us to draw upon a large body of results [7, 8, 22, 30].

Since *A* is stable, the community reaches a unique, globally-stable equilibrium, and the sub-community of coexisting species is characterized by a feasibility and non-invasibility condition [20]. Importantly, in this case one can prove that the effect of the mean interaction strength *μ* on the resulting community is relatively straightforward: *μ* does not affect the identity of the coexisting species, and rescales their biomasses by a constant (see supplementary information, section S6, for details). Similarly, any choice of *r* > 0 only rescales the equilibrium biomasses. Thus, without loss of generality, we can assume *μ* = 0, and *r* = 1, so that the regional pool is completely characterized by *A*.

To describe the statistical properties of the community of coexisting species, we need to condition the distribution of the variables of interest on the *unique* feasible and non-invasible sub-community for a given species pool 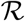. We focus on the following properties: the distribution of the number of coexisting species, the total biomass of the community, and the relative abundance distribution of the coexisting species.

To illustrate our results with an empirical tree structure, we take the phylogeny of the clade *Senna* (Fabales) as an example [39]. The tree contains a total of 94 species and we use the outlier group comprising the species *Senna silvestris var guarantica, Senna siamea, Senna polyantha* and *Senna galeottiana* to root the subtree containing the remaining 90 species.

Notice that, as shown by [32] the final community composition reached in each of our theoretical experiments is the same as would be reached under sequential, one-at-a-time species invasions. Thus, our results map directly to the process of bottom-up assembly of ecological communities. In particular, our results can be used to infer properties of the assembly graph 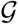 associated with each regional pool [12].

## Results

### Deterministic Limit

First, consider the case where the number of traits, *ℓ*, is very large relative to the number of species, *n*. Let 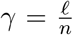 be their ratio. In the limit *γ* → ∞ we find that *A* → ∑ (i.e., the sample covariance matrix converges to the population covariance matrix). Thus, the properties of the community are determined solely by ∑. The simplest case to study is if ∑ = *I_n_* (the identity matrix), which represents the covariance matrix induced by the degenerate *n*-star tree with 0 shared branch length among all species (see Figure 2). This covariance structure corresponds to an evolutionary scenario where all species diverge immediately at time 0. In this case, complete coexistence of 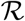 and any of its sub-communities 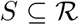 follows trivially, since species do not interact with each other. Remarkably, the same behavior (full coexistence) is shared by any ∑ induced by a tree *T*. This can be proved inductively using the following observation: If *t*_1_ is the time at which the first split happens in the phylogenetic tree, then “cutting” the tree at this branching point generates non-interacting sub-trees *T_i_*, which we assume to have fully coexisting equilibria under the induction hypothesis. Pasting these sub-trees together at their roots gives us a degenerate tree for which the induced covariance matrix is a block diagonal matrix 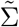. This operation preserves coexistence, since the sub-trees are still non-interacting. We recover *T* by attaching a branch length *t*_1_ to the root. In terms of the variance-covariance matrix, ∑ is obtained by adding a constant to 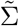. Assuming equal growth rates, this transformation does not affect the feasibility of the system; hence ∑ has a feasible equilibrium (see fig S1 and supplementary information S2 for a more detailed argument). Thus, for *γ* ≫ 1 (i.e., if *A* ≈ ∑), we have full coexistence regardless of the tree topology. Moreover, the assembly graph 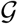 for the species pool 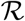 contains all possible assembly histories (c.f. [32] and [42]); in other words, any sub-community can be built by starting with a single species and adding the remaining members sequentially in any order.

### Total biomass and abundance distribution

Under our model, phylogeny strongly influences the biomass and relative abundance distribution of a community. As illustrative examples, consider the two extreme tree topologies given by the “perfectly unbalanced” tree and the “perfectly balanced” tree (Figure 2). Assuming equal inter-branching times, the total biomass of the system, *W*(*n*), for a pool of *n* species is given by 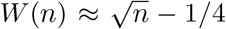 in the perfectly unbalanced case, and 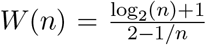 in the perfectly balanced case (see section S2 for details). Similarly, we are able to derive expressions for the individual biomass of each species *s_i_*, where the index corresponds to the position in the ordered tips of the tree (see Figure 2). For the perfectly balanced case, the abundance distribution is trivial, since each species necessarily has the same abundance. On the other hand, the hierarchical nature of the perfectly unbalanced tree is reflected in the individual biomasses, with species that split from the rest early on having much higher abundances. Fig 3 and S2 show that the results are qualitatively unchanged if we sample the inter-branching times from appropriately normalized exponential or uniform distributions. The uneven distribution of abundances for the unbalanced tree helps explain the difference in total biomass: in the perfectly unbalanced case, as *n* grows there is a fraction of species (outliers) that interact less and less strongly with the rest of the community, so that their abundance approaches the limit 1 (obtained for non-interacting species). In contrast, in the perfectly balanced case the abundance of all species is the same, and approximately 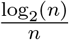.

**Figure 3:**
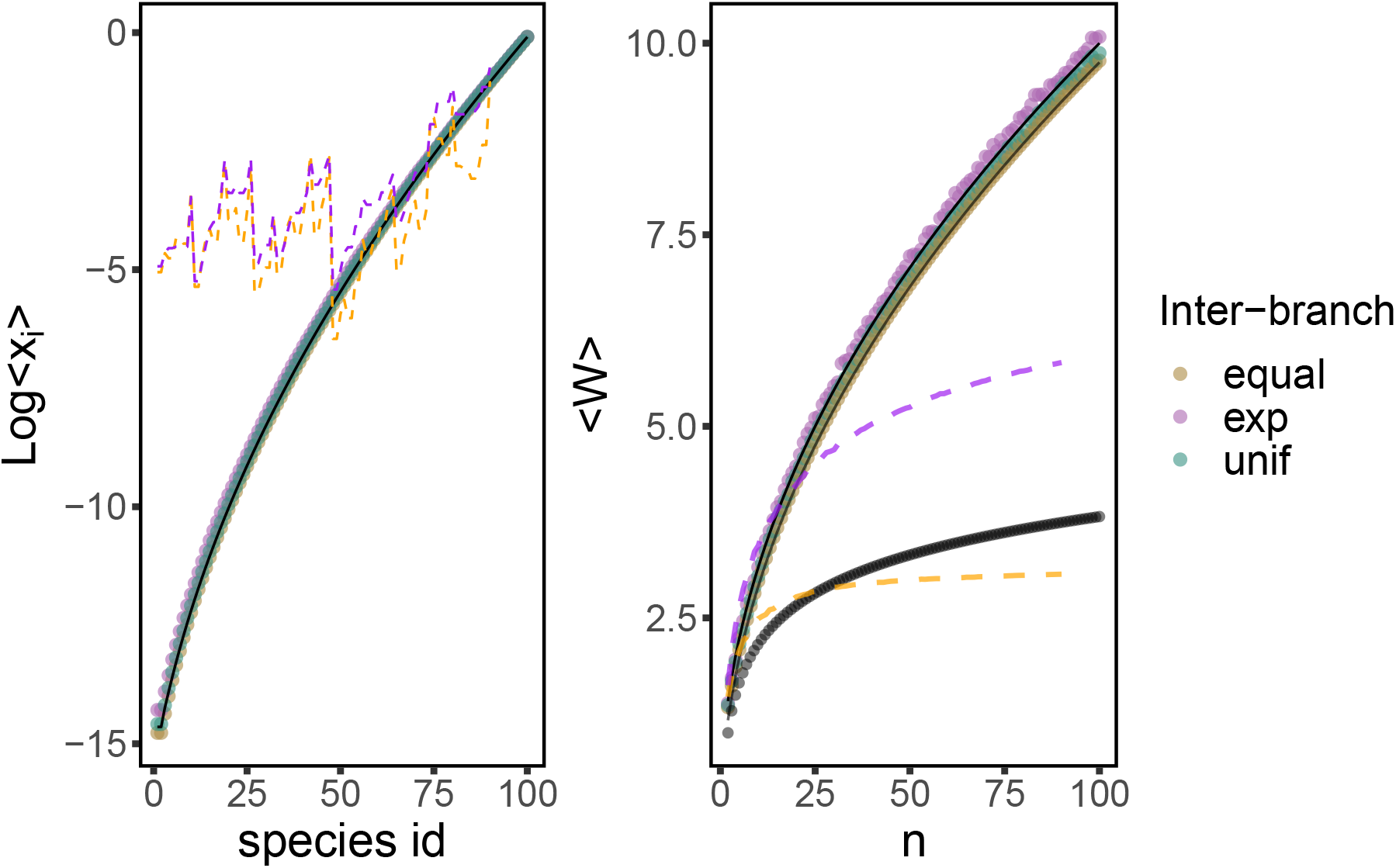
Individual and total abundance for the deterministic limit. Log individual abundance (left) and total abundance (right) for the communities in the deterministic limit of a perfectly unbalanced tree. Dots mark the average values when sampling the branch lengths from an exponential distribution with rate 1, a uniform [0, 1] distribution, and the case of equal branch lengths. The total branch length is renormalized to 1 in all cases. Solid lines are the analytic predictions under equal branch length 1/*n*. In the right panel, the two solid lines are given by 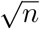 and 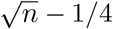, the black dots represent the analytic formula for a perfectly balanced tree which shows logarithmic growth 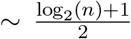. The dashed lines on the right panel are the scaling with size of sub-trees of the *Senna* phylogenetic tree, and the dashed lines on the left plot are the abundance distribution for the full tree (compare with Figure 6). In both cases the purple line is for the case of equal inter-branching times and the orange includes the branch length information.

To compare these results with predicted abundances using a more complicated tree structure, we repeated the calculation using the phylogenetic tree of the *Senna* clade (Fabales) [39]. We considered two cases: either we include the branch length information, or we set all branch lengths to be equal, so that only the effects due to the shape of the tree are considered. The average total biomass *W*(*n*) for sub-communities of different sizes (Figure 3) shows that for both cases at small enough sizes, *W*(*n*) behaves as predicted by the perfectly unbalanced model—which reflects the hierarchical low-level structure of the tree (Figure 6). But as the size of the sub-community increases, *W*(*n*) either reaches values even smaller than the perfectly unbalanced tree, or settles in the middle of the two—showing that, under equal inter-branching times, the perfectly balanced and perfectly unbalanced tree represent the two extreme topologies. The species’ abundance distribution, as for the perfectly unbalanced tree, reflects the tree structure: the abundance profile shows peaks at each of the outliers within clades, and an overall decreasing trend toward more deeply nested parts of the tree (see also Figure 6).

### Star phylogenies

Classical results in theoretical ecology have extended the principle of competitive exclusion to the case of multiple resources/regulating factors, showing that a necessary condition to observe a non-degenerate coexisting community of *n* species in our model is *ℓ ≥ n* [24, 41]. We have shown above that, for a fixed size of the pool, *n*, coexistence is guaranteed in the *ℓ* → ∞. To characterize the cases in between *ℓ = n* and *ℓ* → ∞, we exploit the fact that A follows the Wishart distribution; as such we can make use of tools developed in statistics and economics to explore how the limit of full coexistence is approached (see section S3). To start, let ∑ be induced by a star-tree with shared root of length *ρ* (see Figure 1). In this setting, there is a constant correlation *ρ* among the species in the pool. We find that for *γ* ≈ 1, full coexistence is never achieved for large enough communities (Fig S3). Nevertheless, the community does not collapse completely, and a non-vanishing fraction of species is observed to coexist (Figure 4). The effect of increasing correlation among species is, as expected, to reduce the proportion of coexisting species, *ρ*. In particular, to observe at least half of the species to coexist (in expectation), we need:

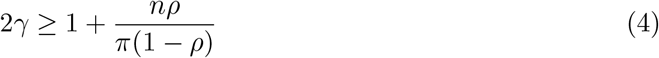

**Figure 4:**
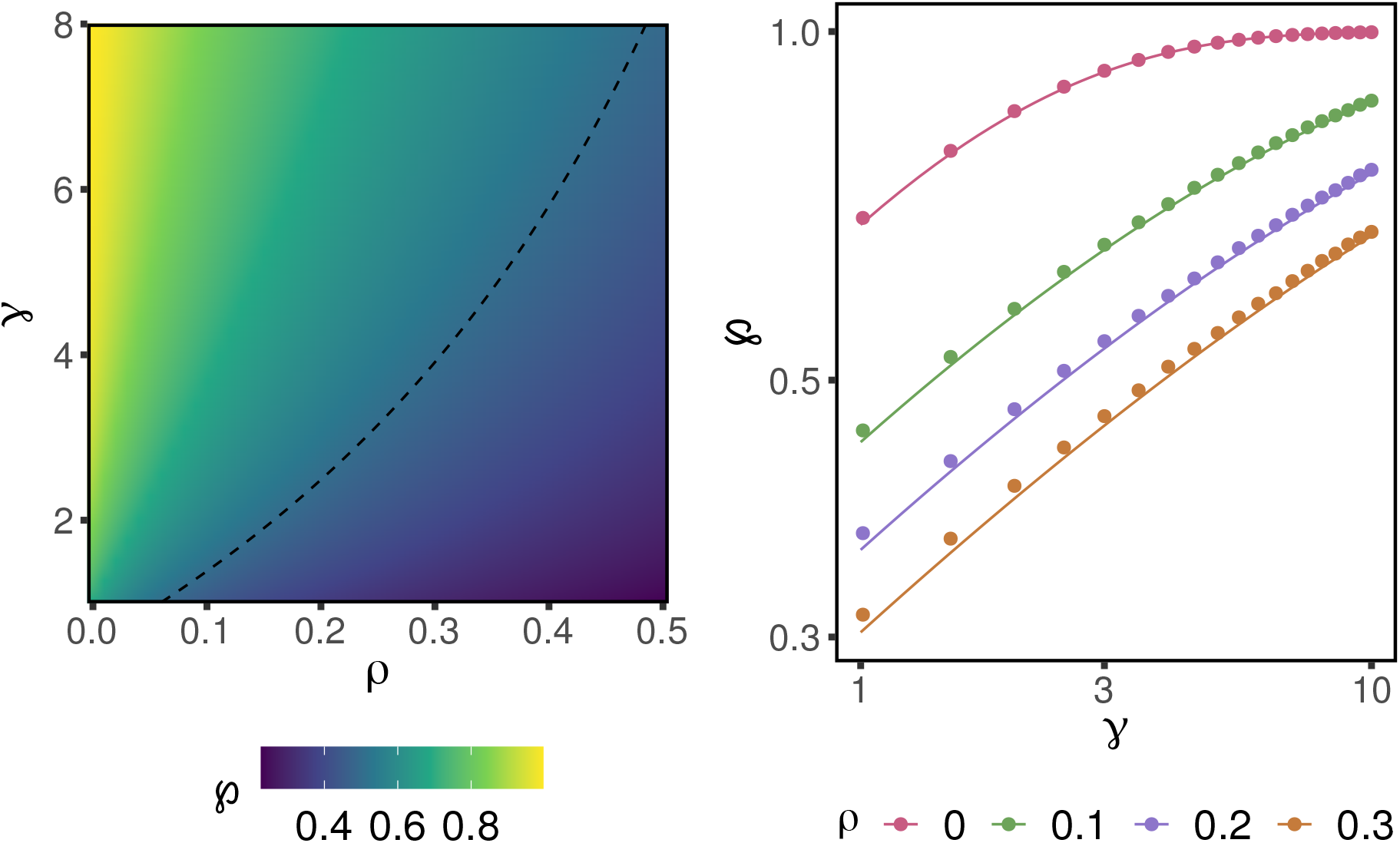
**Proportion of coexisting species 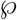 as a function of *ρ* and *γ*.** In the right panel, we compare our analytical approximations (solid lines) with simulations (dots) for a regional pool of 50 species (log-log scale). The left panel explores in more detail the parameter space (*γ, ρ*). The dashed line marks parameters for which we expect half of the species to coexist. As suggested by eq. (4), the value of *γ* giving a fixed 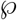 increases sharply with *ρ*.

The quantity 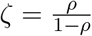 could be interpreted in the framework of population genetics as the ratio of shared to private mutations for each species. It is a key quantity, in the sense that two distinct pools 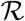 and 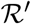, of sizes *n* and *n′* will yield the same mean fraction of coexisting species, for a given *γ*, whenever *nζ = n′ζ′*.

The distribution of total biomass, *W*, for the community of coexisting species is influenced by *γ* and *ρ* in two different ways: the parameters affect both the distribution of the number of coexisting species, and the conditional distribution of *W* for a given community size. Assuming that the distribution of the number of coexisting species is highly peaked at the mode, we derive an approximation for the mean of *W* that closely matches results from simulations (see section S4 and figure S5 for exact results and full distribution). For small enough *γ* the variance of the interactions allows the possibility of positive interactions that enhance *W*, as *γ* increases the interaction matrix converge to the purely competitive interaction matrix given by ∑. This convergence explains the decrease of *W* with *γ* depicted in Figure 5.

**Figure 5:**
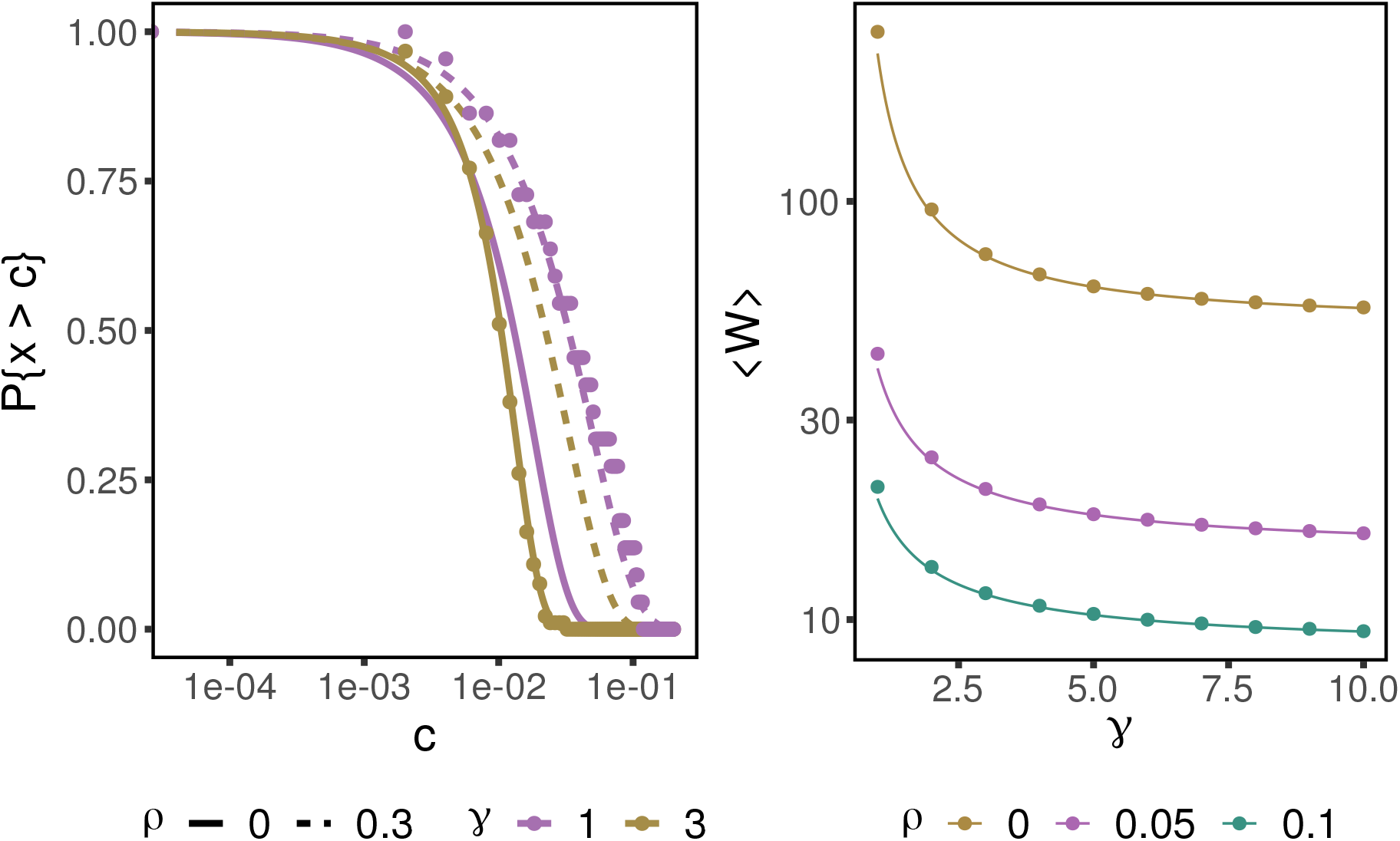
Mean total biomass and relative abundance distribution. The panel on the right shows (note the log-transformation for the y-axis) the mean total biomass for the community of coexisting species; the points represent simulations, and solid lines the corresponding analytical approximations for a pool of 50 species (see section S6 for the effect of changing *μ*). The total biomass decreases as *γ* grows, because the overall strength of interaction between species decreases. The survival function for the relative abundance values of the community is plotted on the left panel (note the log x-axis), where again points stand for simulations and lines for analytical predictions for distinct *γ* and *ρ* values, and a pool of 100 species. For clarity, we just show simulations for the parameters (*ρ, γ*) ∈ {(0,3), (0.3,1)}. In particular we have that as *γ* increases the distribution becomes more and more peaked (as expected) while increasing *ρ* flattens the distribution.

Using the same strategy, we are able to derive approximations (see section S5 for exact formula) for the survival function of the relative abundance distribution under distinct values of *ρ* and *γ*. In particular, the distribution becomes very peaked as *γ* increases, while increasing *ρ* tends to make the distribution flatter (Figure 5).

### Beyond constant correlation

Imposing a more general covariance structure ∑ is challenging from a mathematical standpoint, due to the breaking of the statistical equivalence among species—species in distinct parts of the tree have now different statistical properties. In the general case, the identities of the species matter, and instead of looking at the total number of coexisting species, we focus on how the probability that a particular species survives (*p_s_*) changes with its position in the tree. Simulations for the phylogenetic tree of the *Senna* clade show that the model recreates the phenomenon of phylogenetic over-dispersion: for a group of closely related species, *p_s_* peaks at the outliers of the clade. Furthermore *p_s_* reflects the tree structure in the same manner as the total abundance distribution (compare Figures 1 and 6).

**Figure 6:**
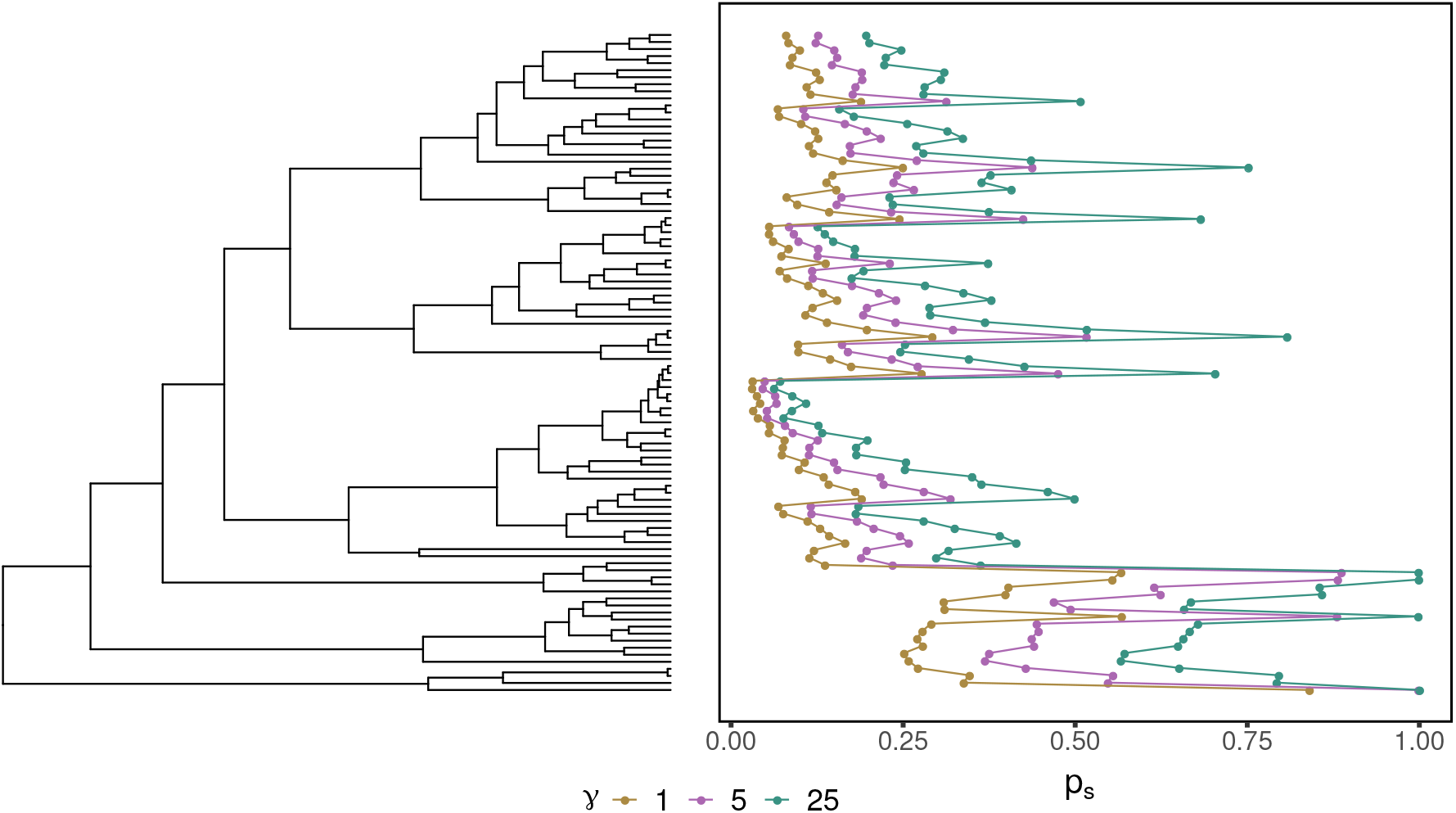
Probability of individual species survival for an empirical tree. The probability that a species is observed in the community of coexisting species, *p_s_*, out of 5000 simulations, is shown alongside the phylogenetic tree (*Senna* clade) where the outermost group is used to set the root. The values *p_s_* reflects the tree structure and the abundance distribution showed in Figure 1: The peaks in *p_s_* correspond to outliers within a group of closely related species and *p_s_* has a decreasing trend towards the most nested parts of the tree (upward direction). In particular, the model produces phylogenetic over-dispersion.

To further explore this relationship, we are able to analytically compute the probability of observing each sub-community in a three-species community (Figure 7). For *n* = 3, there is only one possible tree topology, and we consider the case where all branch lengths are equal. We find that sub-communities containing the outlier species, *s*_3_, are always more likely to be observed than sub-communities of the same size in which S3 is absent. More generally, the formulas in section S3 can be evaluated numerically to find the probability of observing a particular sub-community under an arbitrary, not necessarily ultrametric, tree structure.

**Figure 7:**
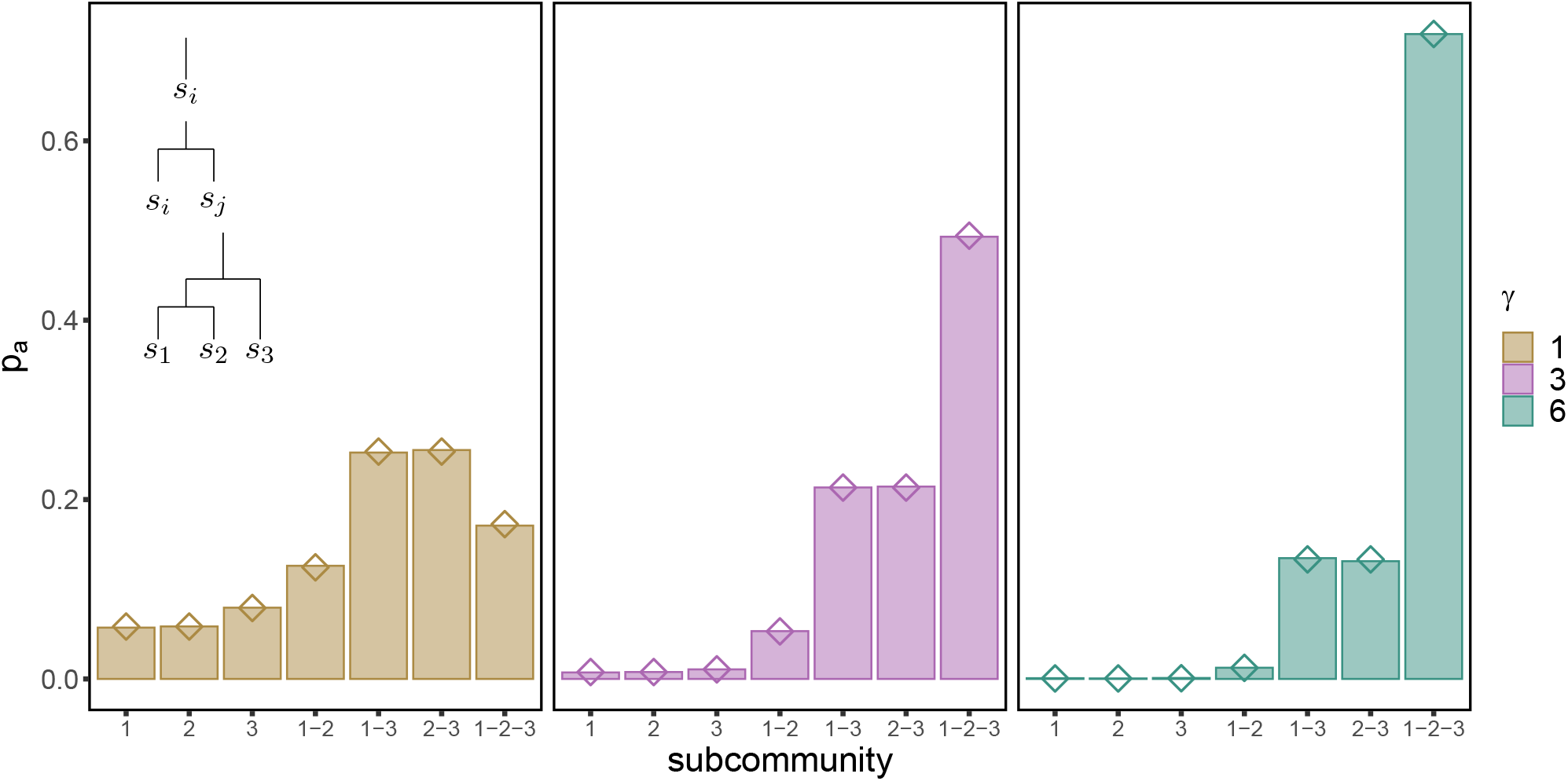
Sub-communities of perfectly unbalanced tree. Probability of observing a given sub-community of the three-species tree with equal branch lengths. The inset shows the tree sub-structures for each of the sub-communities. Bars represent frequencies over 50000 simulations, and dots the analytical predictions.

## Discussion

By considering local community dynamics in a trait-based interaction model, our results provide a clear link between the phylogeny of the regional species pool and many aspects of species coexistence. Importantly, while the tree structure is reflected in the local community patterns, the number of traits controlling interspecific interactions modulates the outcomes.

We found that, when phylogenetic relatedness completely controls interactions, i.e., when the number of traits is sufficiently high compared to the number of species, full coexistence of any sub-community is guaranteed. This result requires both the tree structure (which induces a particular interaction matrix) and the assumption that all species have equal growth rates.

While we expect this result to hold qualitatively for small deviations from the assumption of identical growth rates (see section S7), it is false in general when these requirements are not satisfied. Under the same assumptions, the abundance distribution of the community reflects the tree structure at distinct levels: high biomass is observed for the outliers within each clade (local tree structure), and one expects an overall decreasing trend towards more nested parts of the tree (coarser structure).

When the number of traits is comparable to the number of species, our model is an instance of a Lotka-Volterra model with random interactions. The analysis of models considering random interactions between species has a long tradition in ecology [1, 17, 27], and in recent years the field has moved beyond questions concerning the stability and feasibility of the whole system, focusing more closely on the properties of sub-communities that coexist through the dynamics [3, 6, 10, 33]. In these models, one must usually assume that species interactions are independent of species identity (but see [2, 16]). The star-tree case studied above satisfies this assumption, but with a stronger correlation structure than has been previously considered. This case behaves much like other random interaction models: full coexistence of many species is extremely unlikely, but we expect a moderate number of species to coexist. We have derived an approximation for the mean number of coexisting species, which depends on the ratio of traits to species, *γ*, and on the expected ratio of shared to private mutations for each species *nζ*. As long as these two quantities are the same, pools of distinct sizes will yield the same distribution for the number of coexisting species. Contrary to previous studies [33], the analytic tractability of the model allows us to derive exact expressions for the total biomass and relative abundance distribution of the system.

The general case of an arbitrary, tree-induced correlation structure provides a biologically-meaningful way to relax the statistical equivalence between species. Taking advantage of the vast literature on the Wishart ensemble in fields ranging from economics to statistics [8, 22, 30], we are able to derive exact integral formulas to compute the probability of survival for any sub-community under arbitrary tree structure. In this way, one can measure properties of the system (conditioning on a final sub-community) by numerically evaluating the integral expression. For small enough communities and simple enough phylogenies, this approach can be replicated on each sub-system to compute the marginal distribution of the properties of interest. However, as the number of species grows, these calculations become burdensome. As such, devising new analytical techniques to tackle the general case would be an important step toward studying more general random interaction models, and also advance our understanding of the effects of phylogenies on communities.

Our approach can be extended in a variety of ways, and we briefly discuss some of the most promising avenues.

First, instead of assuming that the same tree structure controls the evolution of all *ℓ* traits, we can partition the traits into *m* classes and assume that the evolution of each class is determined by a distinct phylogenetic tree. These type of processes are studied in population genetics when either admixture or incomplete lineage sorting lead to traits that cannot be explained by a single tree [31]. In such cases, A would no longer follow the Wishart distribution but would rather be a sum of (possibly degenerate) Wishart matrices.

Second, our assumption of equal growth rates among species allowed us to examine how phylogenetic relatedness influences coexistence in a purely interaction-driven model. When we include variation in growth rates, we expect our results to hold only for sufficiently small variance. In this case, the restriction on *l ≥ n* can be lifted, provided that there is a background competitive effect *μ* strong enough to prevent divergence of the dynamics (see section S7). It would be interesting to consider models where growth rates vary under the influence of phylogeny; by modulating how strongly evolutionary relatedness affects both growth rates and interactions, one could investigate the duality between “competition” and “filtering” that is frequently discussed in the literature [14, 28, 38].

Lastly, our approach assumes an explicit separation between evolutionary processes at the regional level (which give rise to the phylogenetic structure) and ecological interactions (at the local level). To remove this separation, one could model the tree generation process and ecological dynamics concurrently. For example, as done by Maynard et al. [29], one could “run” the dynamics after each new speciation event, thereby pruning the community to a coexisting sub-community. One would then take the sub-tree of that community as the starting point for the new speciation event. In this setting, in a similar manner to studies of community assembly [32] and the framework of adaptive dynamics [21], we have a separation of time-scales between the speciation events and the local community dynamics. Traits evolve on the tree between each pruning event. In this regard, our results provide baseline comparisons, and even suggests patterns that would emerge from the process: assuming that the number of traits is a constant *ℓ*, the community cannot reach more than *ℓ* species, yet at the early steps of the process the ratio of traits to species could be extremely high—hence we expect that most speciation events occurring early on would not cause extinctions; in this case, the bulk of the phylogenetic structure would be built at the beginning of the process. Perturbing the growth rates slightly, one could compare the structure of this tree with the structure of the tree found by simply letting the tree generation process run, and after having the same number of speciation events let the species interact and get a coexisting sub-community.

While there has been extensive discussion of the potential ways in which phylogeny could affect ecological differences, and thus interactions, among species [11], much less has been said about the patterns one would observe under a particular hypothesis. In this work, by linking phylogenies to a simple model of trait evolution and local community dynamics, we were able to fully characterize many global aspects of the community. We showed that the phylogenetic structure of the species pool and the the number of traits determining competition affect the results in concert. Our results provide a useful baseline prediction for the effect of phylogeny on community dynamics and coexistence.

## Acknowledgments

We thank P. Lemos and M.O. Carlson for comments on the manuscript. J.A.C. acknowledges financial support from the Spanish ‘Ministerio de Economía y Competitividad’ project PGC2018-096577-B-I00. Z.R.M. acknowledges support from the National Science Foundation Graduate Research Fellowship Program under Grant No.(DGE-1746045).

## Supplementary information

### 1 Motivation

#### From consumer-resource dynamics to covariances

We start with a model of consumer-resource dynamics in which the consumers differ only in the relative preference of each resource and the resources have an homogenous growth rate. Let 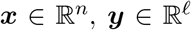 be vectors denoting the density of predators and resources. We model the dynamics as the MacArthur’s consumer-resource model [25]:

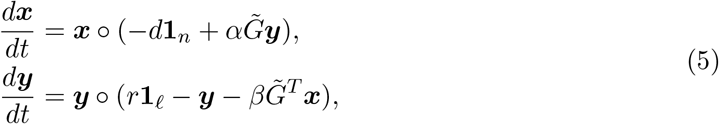

where ○ stands for the Hadamard (component-wise) matrix product, and 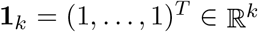 is a notation for a column vector whose entries are exactly *k* ones.

By our assumptions, matrix 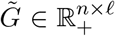 encodes the preference distribution (alternatively, the time allocation distribution) of the predators over the resources, so that 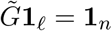. Then by a separation of time scales, which implies that resource densities remain at equilibrium, we can model the competition between the consumers as following competitive Lotka-Volterra dynamics [25]:

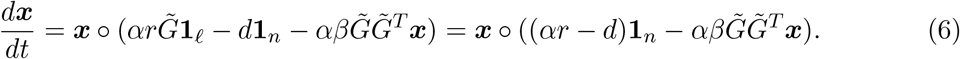

As long as *n ≤ ℓ* (besides measure zero sets) we have that matrix 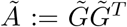 is positive definite. This property of *Ã* allows one to further transform the system (6) without affecting the set of coexisting species. In particular we can perform the following operations (see section 6 for a more detailed discussion):

a. Rescale the growth rate, ***υ*** = (*αr − d*)**1**_*n*_, by any positive constant.
b. Multiply *Ã* by a positive, constant diagonal matrix.
c. Multiply both *Ã* and *υ* by a positive diagonal matrix.

Following this operations we reduce the system to

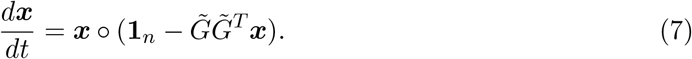

To distinguish the effect of the mean of 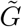, write 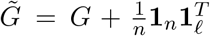. Notice that this decomposition, together with the restriction 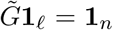, implies that *G***1**_*ℓ*_ = **0**_*n*_, which means that the entries of *G* have zero mean —here **0**_*k*_ = (0,…, 0)^*T*^ stands for a column vector formed by *k* zeros. Then matrix *Ã* can be decomposed as 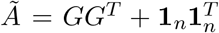. Because the system in (7) has constant growth rates then one can show (section 6) that, as long as *ℓ > n* (the strict inequality arising due to *G* having rank *ℓ* − 1), the set of coexisting species for (7) is invariant to the shift 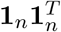. Therefore the system reduces to:

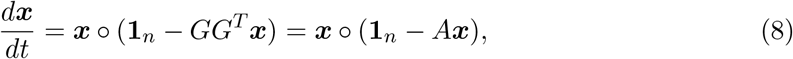

where we have defined *A* := *GG^T^*. This is the competitive, deterministic dynamics that we have assumed for consumers throughout this contribution. Observe that the set of coexisting species remains unchanged if we define interaction matrix 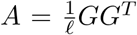, as in the main text, because of the aforementioned invariant operations.

#### Modelling the covariance matrix

From (8) we see that the interactions between species *A_ij_* are fully determined by the row vectors ***G**_i_*. Because each row 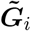 of matrix 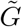 is a preference vector, then it lies on the standard *ℓ* − 1 dimensional simplex 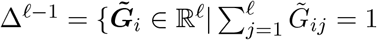, for *i* = 1,…, *n*}, which implies that ***G**_i_* lies on a bounded subset of a linear subspace of 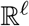 defined by the restrictions 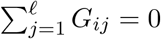 for *i* = 1,…, *n*. By choosing a suitable (linear) coordinate system 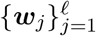 we can express

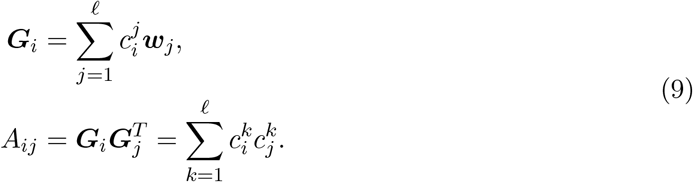

Therefore, the entries of A are fully determined by the coordinates of row vectors ***G**_i_* on the basis 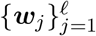.

To model coordinates 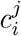 we assume that each (rescaled) preference vector ***G**_i_* is the result of a diffusion process starting at the origin of this space (this maps back to our 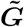 matrix as saying that every consumer has an *homogeneous* preference for any resource). Assuming that each coordinate is independent and letting the diffusion time be small enough, then coefficients 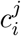 are normally distributed, 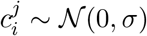. The invariant properties of the model allow us to forget about the deviation *σ* and simply model 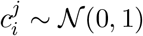. This shows that A satisfies the assumptions of model (8) up to a change of number of traits from *ℓ* to *ℓ* − 1.

### 2 Deterministic limit

#### Full coexistence

We provide more details for the proof that, in the deterministic limit, every subcommunity of the pool is feasible. Since every subcommunity has an interaction matrix induced by a tree, it is enough to show that feasibility is guaranteed whenever this is the case.

We proceed by induction on the number of species. For *n* = 1 the claim holds trivially. Let *T* be a phylogenetic tree (not necessarily ultrametric) for *n* species, and ∑ its respective covariance matrix. Let *t*_1_ be the time at which the first split happens, so that at *t*_1_ the ancestral branch splits into *m* ≥ 2 lineages (*L_i_*, with *i* = 1,…, *m*) where each *L_i_* contains at most *n* − 1 species. Lineages are defined by the condition that species *j, k ∈ L_i_* if and only if the shared branch length between both species *t*(*j, k*) satisfies *t*(*j, k*) > *t*_1_. That is, each lineage contains the subset of species whose shared evolutionary time is strictly greater than *t*_1_. For each *L_i_*, take *T_i_* to be the subtree induced by *L_i_*. Consider 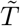, the tree obtained by shrinking the segment between the root and *t*_1_ to a point (see Fig. 8), then 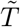 is a phylogenetic tree, for which the covariance matrix 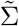 is block diagonal and given by diagonal blocks 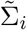. Each 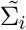 is the covariance matrix of the tree 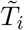 which is obtained from *T_i_* by shrinking the root branch by *t*_1_. By induction it follows that each block 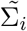 is feasible, hence 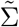 is also feasible. Observe that, going backwards, *T* is obtained from 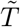 by adding a root segment of length *t*_1_. In particular this says that the shared evolutionary times of all species increases by *t*_1_, i.e. 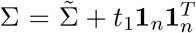, so that ∑ is a constant rank-one update of 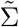. Then by section 6, the equilibrium associated to ∑ is feasible.

**Figure 8:**
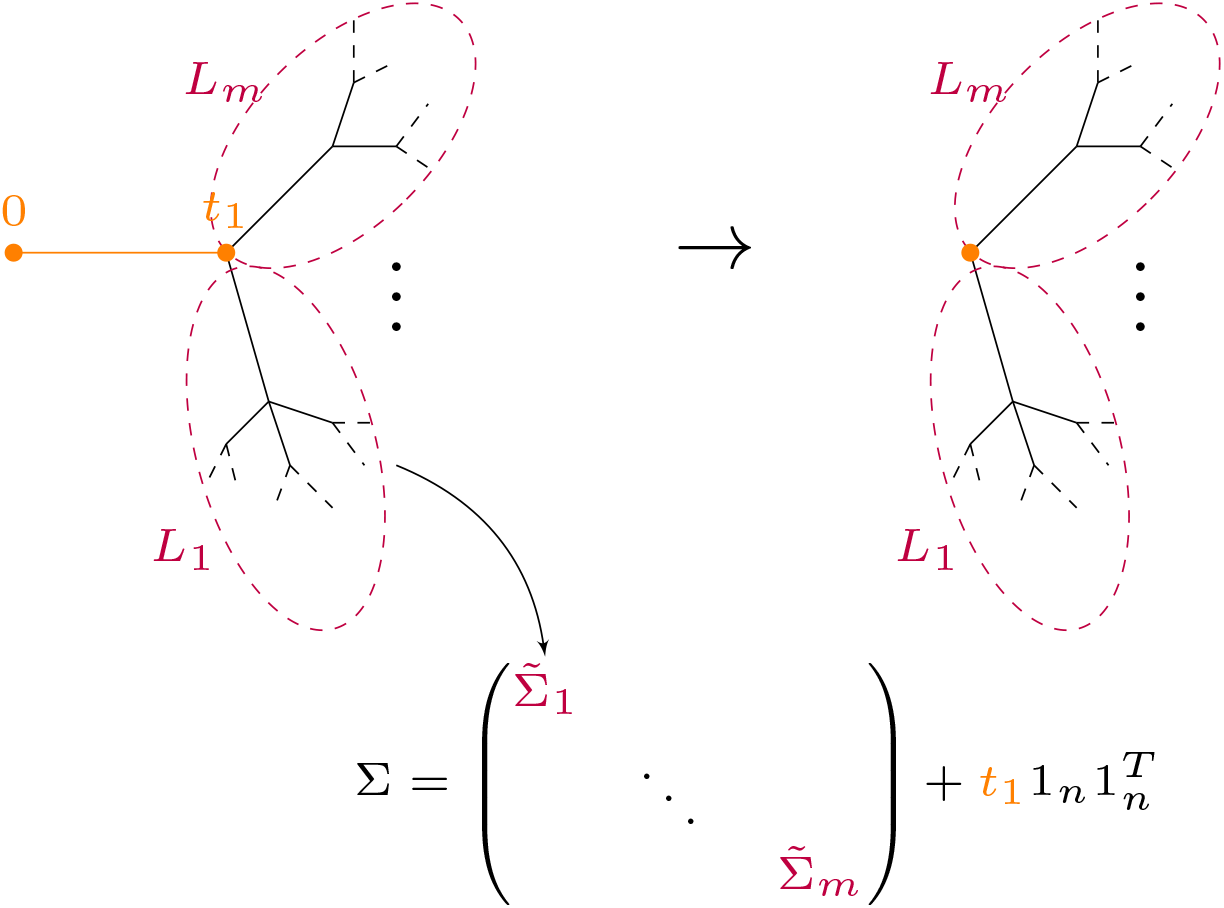
Schematic representation of the inductive step on the proof of full coexistence. Starting with the tree *T* (left), we shrink the ancestral branch up to the first splitting time *t*_1_ to have a degenerate tree 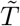 (on the right). 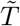 splits at time 0 into *m* distinct subtrees induced by the lineages *L_i_* for *i* = 1,…, *m*. The covariance matrix for *T*, ∑, is obtained from the covariance matrix 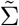 of 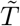 by “adding back” the ancestral branch. This amounts to a constant rank-one update of 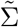 which preserves feasibility.

#### Perfectly hierarchical trees

Consider a perfectly hierarchical tree *T_n_* with *n* tips and branching times *t*_0_ = 0 < *t*_1_ < … < *t_n_* < 1 (see figures 1-2 of the main text), and let ∑_*n*_ be its covariance matrix. Then it follows trivially that

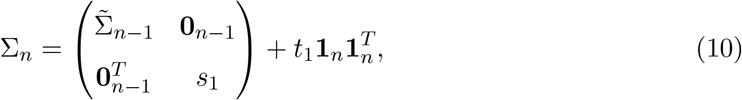

where 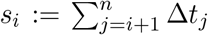, for Δ*t_j_ = t_j_ − t*_*j*−1_ the time between two branching events— the *inter-branching time*. In this subsections we find accurate bounds for the total biomass and analyze the expected abundance distribution.

Define the vector of abundances 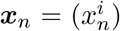 for a hierarchical tree *T_n_* with *n* tips. In the deterministic limit, this vector satisfies the linear system

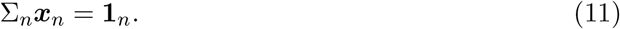

As in the proof of feasibility, ***x**_n_* is given recursively by the updated equilibrium abundances 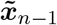 and 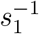 of the non-interacting subtrees 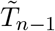 and the one formed by the first species, respectively. Indeed, if we look for solutions of the form 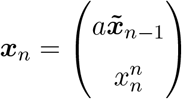, where the vector of abundances 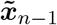 satisfies 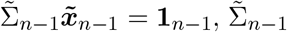 being the covariance matrix of the subtree 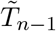, the equilibrium condition (11) for ***x**_n_* reduces to a linear system for *a* and 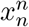:

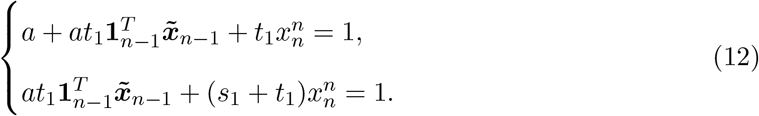

The solution is 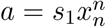, with 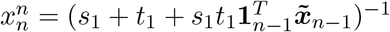. Let 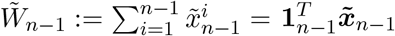. Then ***x**_n_* can be written in terms of 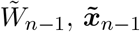, *s*_0_ = *s*_1_ + *t*_1_ and *s*_1_ as

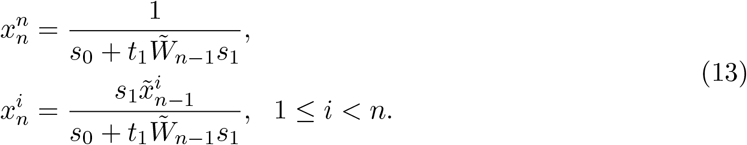

In particular, this implies the following recurrence for the total biomass, *W_n_*:

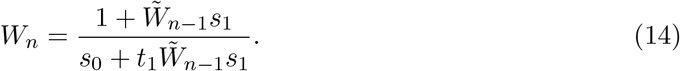

In the case of equal inter-branching times, 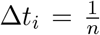 for all *i* = 1,2,…, *n*, observe that *s*_0_ = 1, 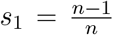 and 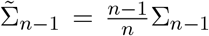. Hence 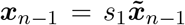 and 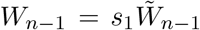, so Eqs. (13) and (14) above reduce to:

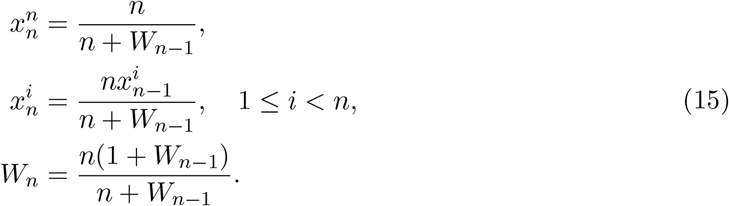

The following proposition provides accurate upper and lower bounds for total biomass in the limit of large number of species.

##### Proposition 1.

*Let*

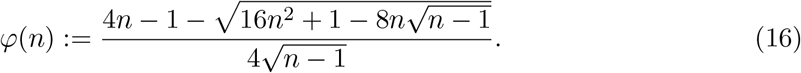

*Then, for equal branching times, it holds that* 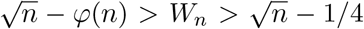 *for n* ≥ 2 *and φ*>(*n*) → 1/4 *in the limit n* → ∞.

*Proof.* Direct computation shows that the inequality holds at *n* = 2 so we proceed by induction on *n*.

Consider first the lower bound. Suppose it holds at *n* − 1, then:

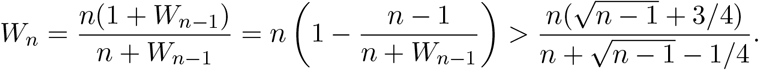

If the claim were not satisfied at *n* we would have

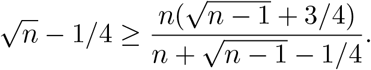

Rearranging terms, this gives the following chain of equivalent inequalities:

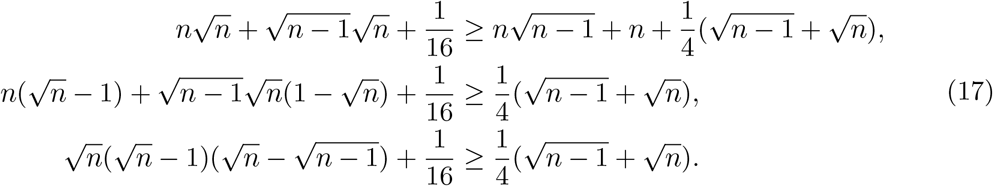

Multiplying both sides by 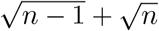 we get

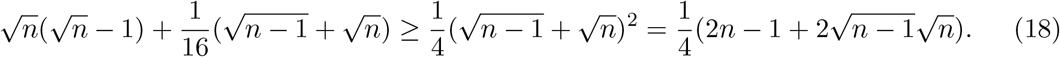

The last inequality implies

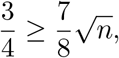

which says *n* ≤ 1. This is a contradiction and we are done.

We proceed in the similar way for the upper bound. By induction hypothesis at *n* − 1 we have

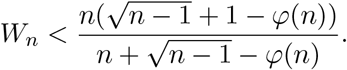

If the inequality is not satisfied at *n* then, a similar chain of inequalities yields

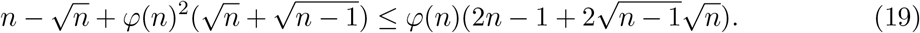

Note that the above restriction is exactly the same as (18) with the inequality reversed and changing *φ*(*n*) instead of 1/4. Using that 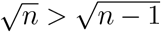, the last inequality implies

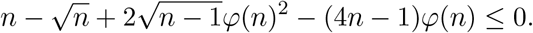

In particular, this means that *φ*(*n*) ≤ *u* for *u* the smaller root of the above quadratic equation,

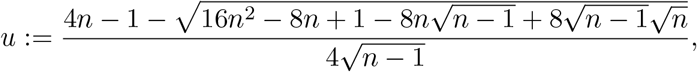

but with this definition and (16) it is easy to see that

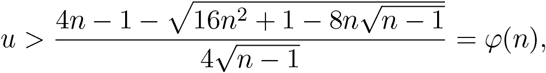

which is again a contradiction and this completes the proof for the upper bound.

We have just proved that 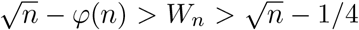. In particular, this implies that *φ*(*n*) < 1/4. Taking the limit in the numerator of expression (16) it is easy to see that the leading order is

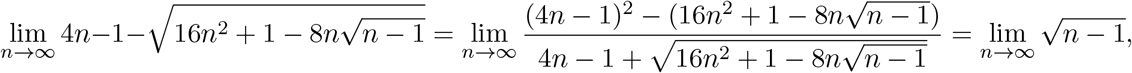

which shows that

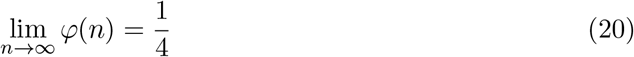

and the proof is complete.

Note that, for large communities, a very good approximation for the total biomass in a perfectly hierarchical tree is given by the formula 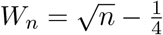.

The recursions in (15) for individual abundances can be easily solved in terms of total biomass *W_n_* as

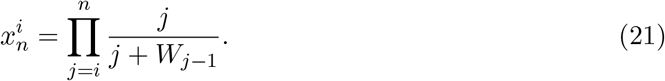

This formula gives the abundance of the *i*-th species (in increasing order of the tips) for *i* ≥ 2 (observe that the first two species have the same abundance). Alternatively,

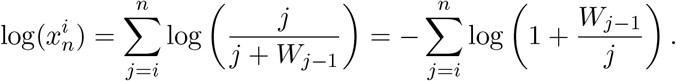

Approximating *W*_*j*−1_ by its lower bound, 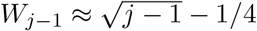, we find

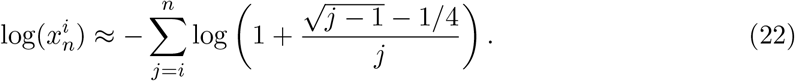

Cutting the series for log(1 + *x*) at second order and considering only the leading term, with respect to *j* for the quadratic term, we get:

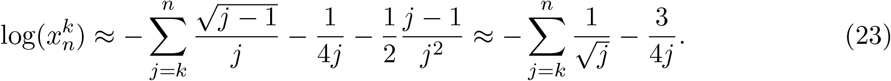

By the Euler-Maclaurin formula we obtain:

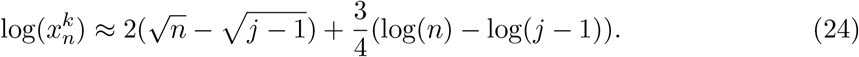

and we can further refine the first terms 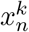 for *k* small by replacing the actual value *W_j_*.

#### Perfectly balanced tree

The total biomass for perfectly balanced trees is easier to derive because the covariance matrix has constant row sums in that case. To show this statement, order tree splits by the time they happen (*t*_1_ < … < *t_q_*). At each time ti, the number of lineages doubles, so we get a total of *n* = 2^*q*^ species. As species interact by their shared evolutionary time, in this case each species shares the time with 2^*q−k*^ other species. Now let 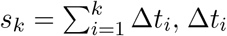 being the inter-branching time —compare the different notation for *s_k_* here and in the previous subsection. Summing over all possible split times we get the sum over any row of A (observe that *A_ii_* = 1),

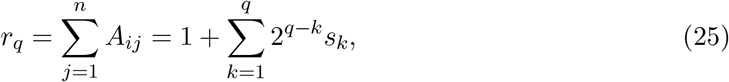

which is independent of *i*. Because row sums are constant, the vector or equilibrium abundances can be written as ***x**_n_ = x***1**_*n*_, and substitution into ∑*_n_**x**_n_* = **1**_*n*_ yields *r_q_x* = 1. Therefore, individual abundances at equilibrium are constant and given by 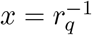. Consequently, the total biomass at equilibrium, *W_q_*, is simply given by

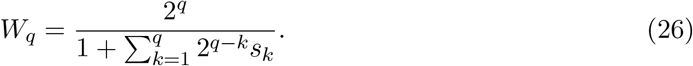

By our assumption of ultrametric trees, we have *s_k_* < 1 (we need to add the tip lengths to sum up to one). In the particular case of equal inter-branching times, 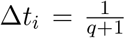, then 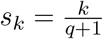 and

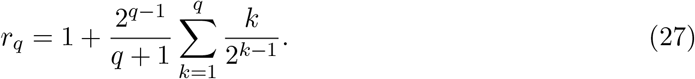

Observe that

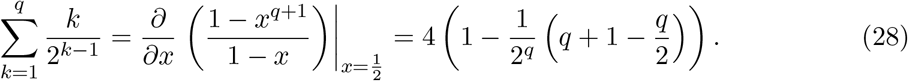

Thus,

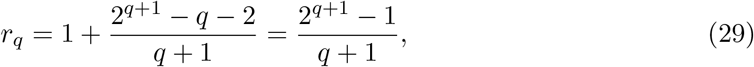

and the total biomass reads

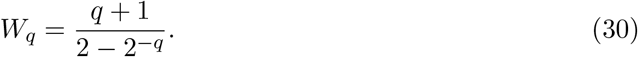

Let *n* = **2**^*q*^ be the number of species, then the number of tree splits is *q* = log_2_(*n*). In terms of the number of species, the formula is given by

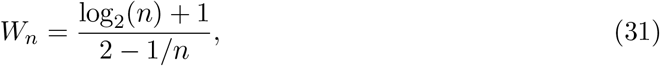

which grows logarithmically with *n*. Fig. 9 compares the case of perfectly balanced tress for equal branching times with two cases, in which sampling times are drawn from exponential and uniform distributions.

**Figure 9:**
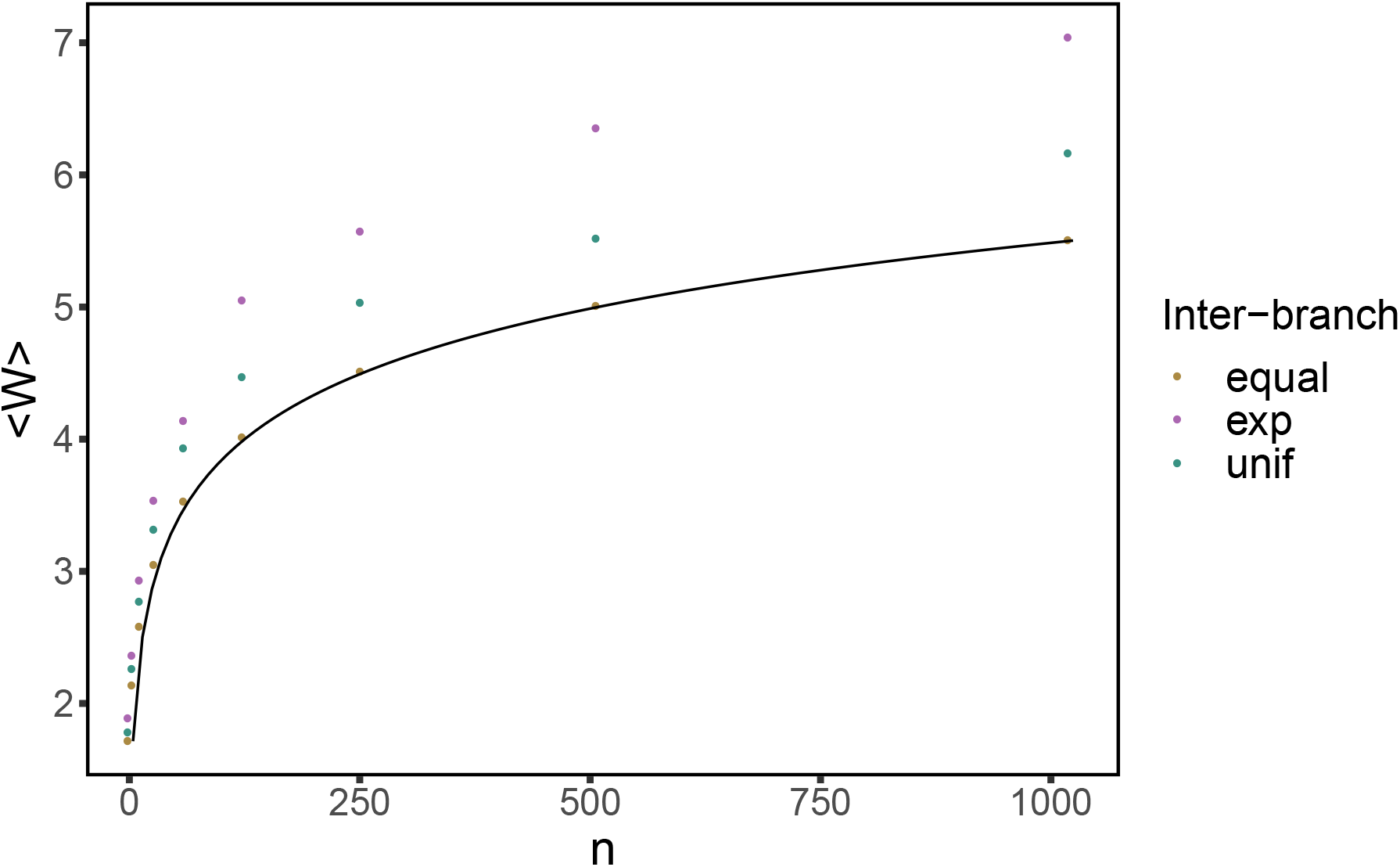
Total biomass for the perfectly balanced tree. Dots mark the average values over simulations when sampling branch lengths from an exponential distribution with rate 1, a uniform [0, 1] distribution, and the case of equal branch lengths, for which the analytical prediction (31) is shown with a solid line.

**Figure 10:**
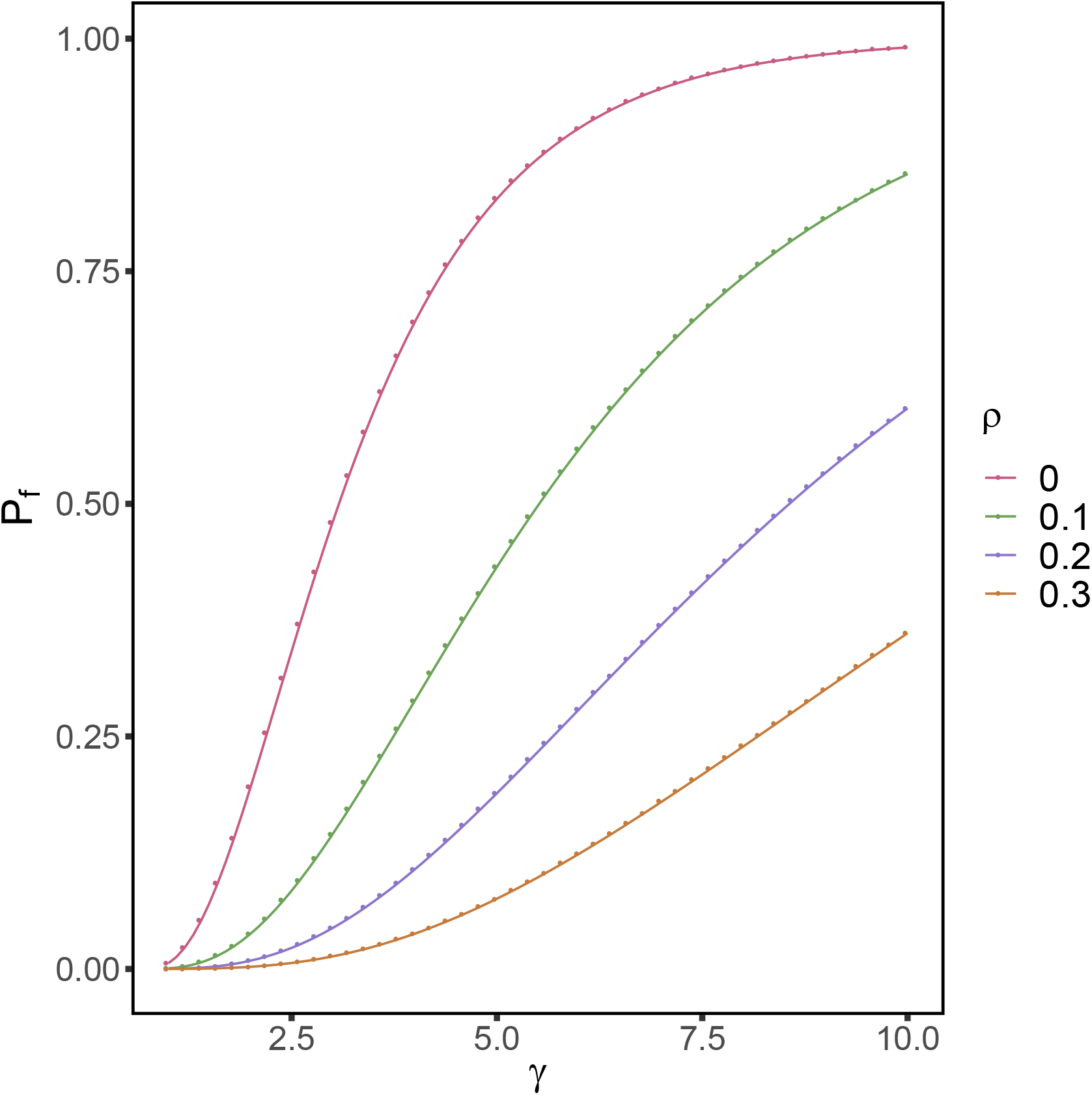
Probability of feasibility as a function of the ratio *γ* of number of traits to number of species for different *constant* correlation matrices. The simulations were done with *n* = 10 species. Dots are simulations, solid lines are numerical evaluations of the exact formula (57). The larger the correlation, the slower curves approach to one in the deterministic limit *γ* → ∞.

**Figure 11:**
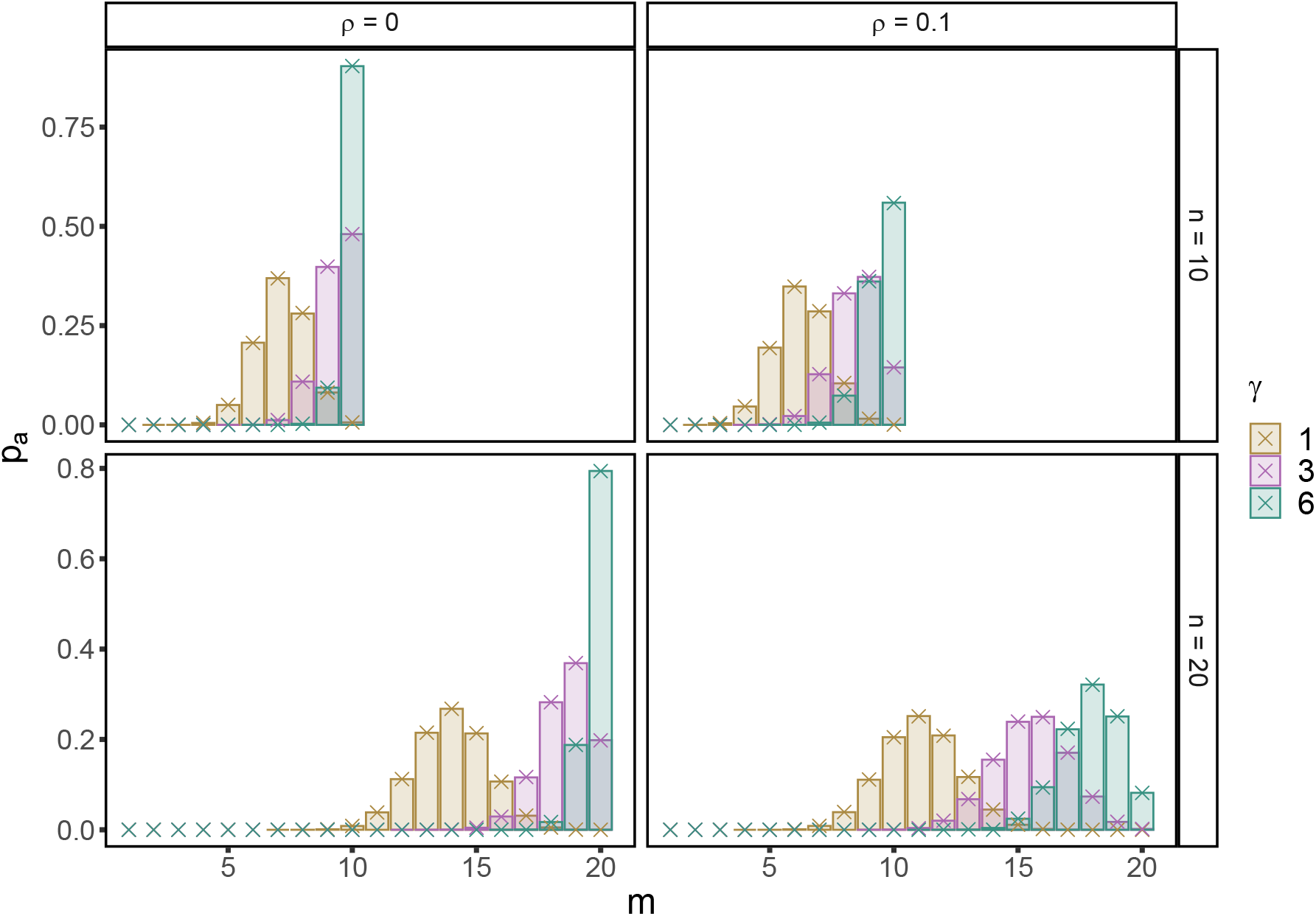
Distribution of the set of coexisting species as a function of the ratio *γ* of number of traits to number of species for different *constant* correlation matrices. The simulations were done with *n* = 10 and 20 species. Bar are simulations, crosses are numerical evaluations of formula (79).

### 3 Number of coexisting species

We have shown above that, in the *ℓ* → ∞ limit, full coexistence is guaranteed. To study species coexistence for finite *ℓ ≥ n* we use the fact that A follows the Wishart distribution. As in [33], first we will compute the probability of the equilibrium point being feasible, i.e., where all species survive. Second, since the attractor is unique (it is the only saturated equilibrium point that appears), we can calculate the probability that the equilibrium point cannot be invaded by the remaining species in the pool. Then we will show that the probability of feasibility and non-invasibility factors into the corresponding product, which yields the distribution of the number of species that coexist, as well as the expected number of species that survive.

Because matrix *A = GG^T^* is symmetric and positive definite, it is diagonally-stable [19], which implies that generalized Lotka-Volterra dynamics exhibits a single, globally stable fixed point [19], so there is a unique endpoint for the dynamics. Let us write the equilibrium abundances of the attractor, formed by *m* survivors, as

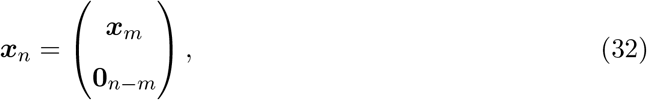

where, without loss of generality, we have located the survivors as the first *m* species. Let {*S*}_*m*_ denote the set of species that survive (i.e., the support of the endpoint). Therefore, the attractor can be fully characterized by two conditions [33]:

- Define the vector 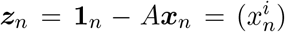 with components 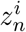. Then it holds: first, 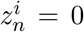 for all species *i* ∈ {*S*}_*m*_, which simply states that equilibrium abundances of survivors satisfy the linear system *A_m_**x**_m_* = **1**_*m*_, for *A_m_* the submatrix of *A* restricted to the support {*S*}_*m*_. Second, it also holds that 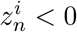 for all species *i* ∉ {*S*}_*m*_, i.e., the fixed point *cannot be invaded* by the remaining species outside the endpoint. We have, therefore, a fixed point that cannot be invaded.
- The equilibrium point hast to be *feasible*, i.e., ***x**_m_* > **0**_*m*_ —here we use the notation that vectors ***a*** > ***b*** if all inequalities are satisfied component-wise.

Since matrix *A* belongs to the Wishart ensemble, these two conditions are to be understood in statistical terms. In the following subsections we are going to compute exact formulae for the probability that all the species in the pool form a *feasible* attractor, and the probability that an endpoint formed by *m* species remains *non-invasible*. Using the properties of the Wishart ensemble [30], we will calculate separately the probabilities of feasibility and non-invasibility, and with them we will obtain the distribution of the number of species that survive.

#### Probability of feasibility

Let *n* be the number of species in the community and *ℓ* the number of traits, and define *γ* := *ℓ/n* as the ratio between the number of traits and the size of the pool. An equilibrium point for the system such that all species coexist satisfies:

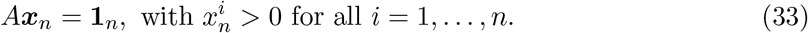

The probability of feasibility is then the probability that *A*^−1^**1**_*n*_ has all entries greater than 0. Observe that interaction matrix is defined as 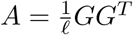 in the main text. Since rescaling by a positive constant in *A* does not affect the condition for feasibility, we can forget about the rescaling by the number of traits *ℓ*.

Let 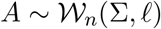 and *L*_*n*−1_ = (*I*_*n*−1_,**0**_*n*−1_) be a rectangular (*n* − 1) × *n* matrix with 0 in its last column, *I_k_* being the *k × k* identity matrix. Then equation (2) of [22] (similarly stated in the proof of Theorem 1 in [8]) implies that

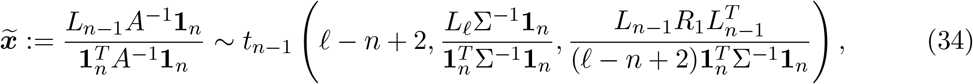

where *t_p_*(*ν, μ*, Λ) is a multivariate, *p*-dimensional *t* distribution with *ν* degrees of freedom, localization vector ***μ*** and dispersion matrix Λ [34]. Matrix *R*_1_ is given by

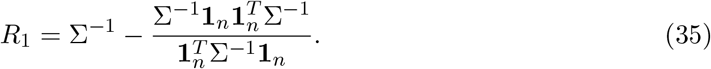

Up to a normalization by a positive constant (which is precisely the total biomass, 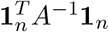, given that *A* is positive definite), vector 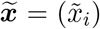 precisely gives the abundances of the *first n* − 1 species. Moreover, the last (normalized) abundance is expressed as 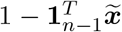, so the probability of feasibility turns out to be

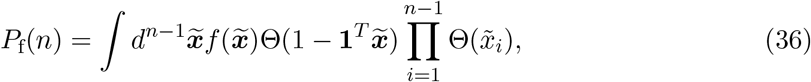

for 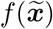 the probability density function of the multivariate *t* distribution defined in (34).

Because a multivariate t distribution is the ratio between a multivariate Gaussian and the square root of a chi-square distribution, it holds that if 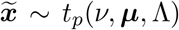, then we have that 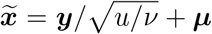, where 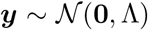 is a multivariate Gaussian and 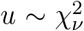, which is independent of ***y***. Therefore, conditioning on *u*, we find that 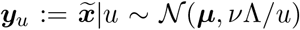 and we can transform the integral above to get

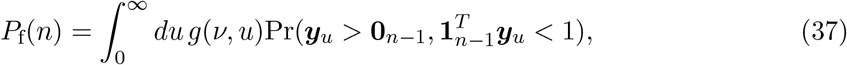

where 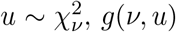, *g*(*ν, u*) is the corresponding pdf with *ν = ℓ − n* + 2, and the random variable ***y**_u_* is distributed as a multivariate normal,

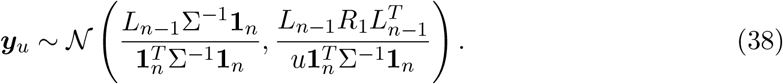

In this way, all the dependence in the number of traits *ℓ* remains included in the chi-square distribution. Eqs. (37) and (38) yield the probability of feasibility for an arbitrary covariance matrix ∑. An explicit calculation of the probability of feasibility amount to evaluating the probability 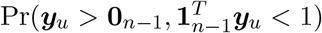. This can be done explicitly for the case of constant, non-negative correlation.

#### Constant, non-negative correlation

Consider the covariance matrix 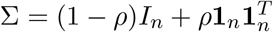 with *ρ* ≥ 0. Then (38) simplifies to:

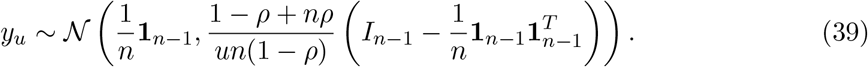

Let us define

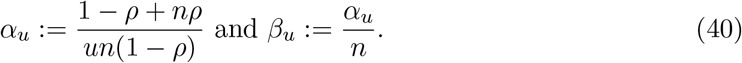

In this way, the covariance matrix ∑_*u*_ in (39) can be expressed as 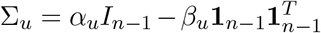. ∑_*u*_ has two eigenvalues, *α_u_* and *α_u_* + (*n* − 1)*β_u_*. The first has multiplicity *n* − 1, and the second 1. Hence the determinant follows immediately,

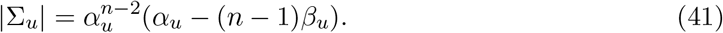

The inverse can be easily calculated:

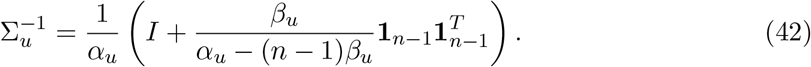

Therefore we can write the pdf for the random variable *y_u_* as

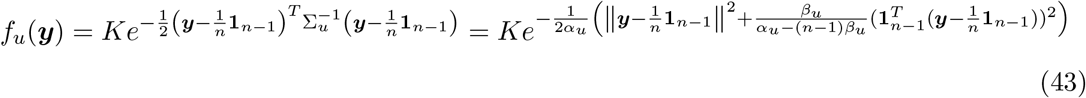

for 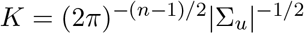. First we have to compute the probability

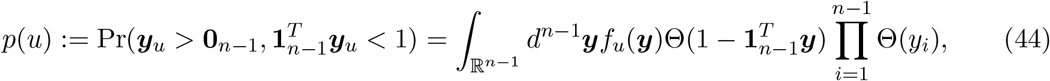

with Θ(*x*) the Heaviside step function, defined as Θ(*x*) = 1 if *x* ≥ 0 and Θ(*x*) = 0 if *x* < 0. Thus after a change of variables 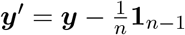, we have

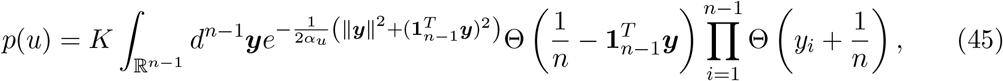

where we have omitted primes to ease notation and we have used (40) to see that

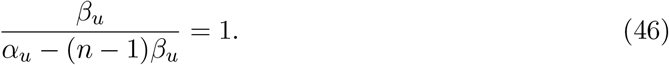

To simplify the term 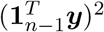 in the exponential, we introduce a Dirac’s delta function,

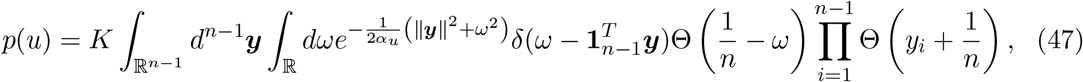

and use its integral representation,

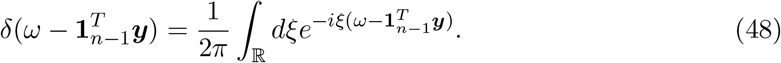

This transformation, together with an interchange in the order of integration, yields the following expression for *p*(*u*):

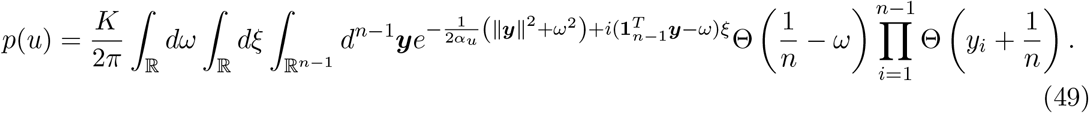

Apparently we are increasing the complexity of the integral, but rearranging terms we observe that

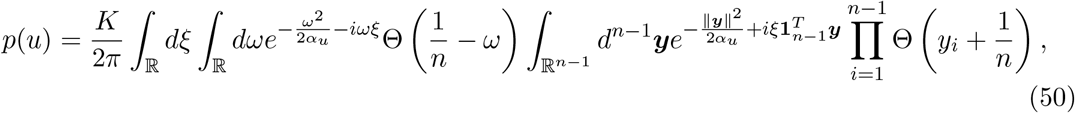

and the integral over ***y*** factorizes,

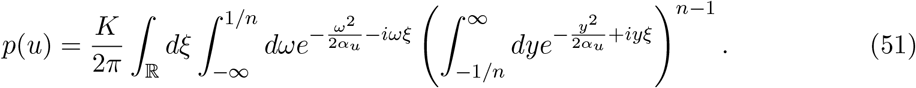

Now, in the integral over *ω*, change to the variable *ω′ = − ω* to get

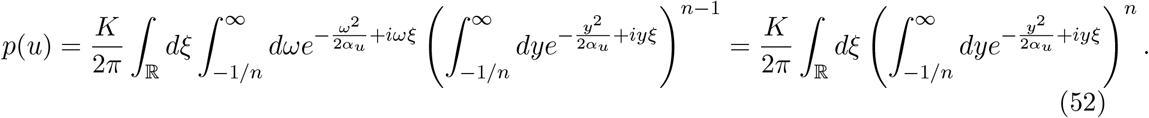

Let

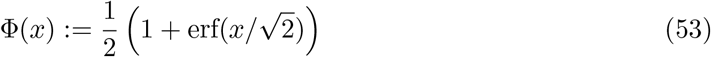

be the cdf of the standard Gaussian distribution, which can be extended to the complex plane.

Then it holds that

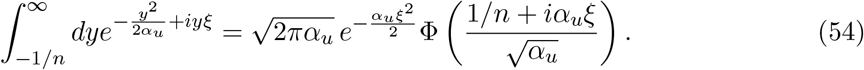

Therefore, the sought probability can be written as

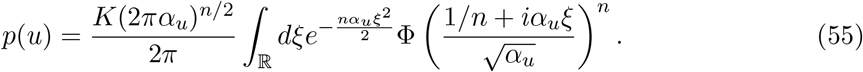

An alternative way to express the integral over *ξ* it is to consider a path Γ in the complex plane such that 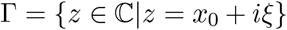 and then reducing the result to the limit *x*_0_ → 0, so that the integral over the imaginary axis is well defined. In practice, this amounts to change to the variable *ζ = iξ*. Consequently, an equivalent form of writing this equation is

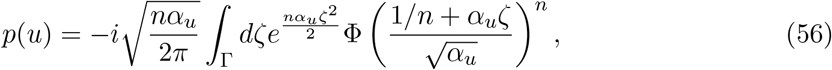

where we have used that 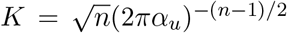 in this case. Finally, according to (37), in the case of constant, positive correlation the probability of feasibility is given by a two dimensional integral,

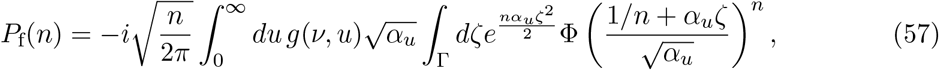

where *g*(*ν, u*) is the pdf of the chi-square distribution with *ν = t − n* + 2 degrees of freedom. Fig. 3 compares this exact formula with numerical simulation for different values of the correlation.

#### Probability of non-invasibility

In this subsection we compute the probability that an attractor formed by *m ≤ n* species cannot be invaded by the remaining *n − m* species. Let *A ~ W_n_*(∑,*ℓ*). Observe that for invasibility the rescaling of interaction matrix as 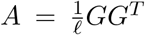 does not matter. Partition matrices *A* and ∑ in four blocks as follows:

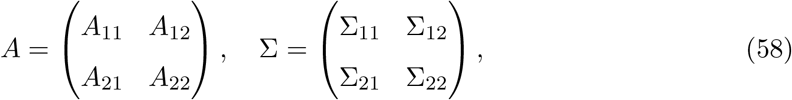

where ∑_11_ refers to the species that belong to the support {*S*}_*m*_ of the attractor, ∑_22_ is related to those species outside the attractor, and off-diagonal matrices are formed by the corresponding rows and columns in {*S*}_*m*_ and {*S*}_*n*_ \ {*S*}_*m*_, and *νice versa*. The exact same notation applies to blocks in *A*.

Then by theorem 3.2.10 of [30] we have that

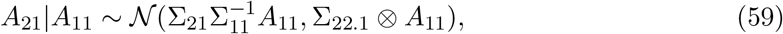

where 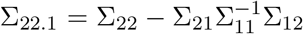 is the Schur complement of ∑_22_, ⊗ is the tensor product of matrices, and the normal distribution appearing is meant to be understood as the distribution of the *flatten* matrix *A*_21_. By the properties of the normal distribution it follows that

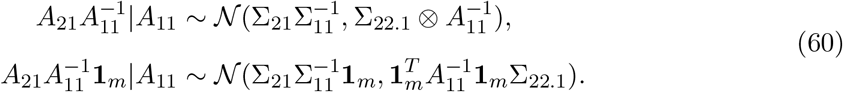

In order to get the last line, we first transpose the matrix, then notice that the 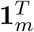 operator acts on the vector of elements of the matrix as *I_m_* ⊗**1**^*T*^. Hence by the property (*A ⊗ B*)(*C ⊗ D*) = *AC ⊗ BD* of the tensor product the second statement above follows.

As mentioned at the begining of Sec. 3, the probability that the attractor cannot be invaded by any species in {*S*}_*n*_ \ {*S*}_*m*_ coincides with the probability that 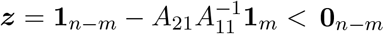. Define 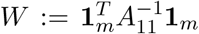 and *f_W_*(*w*) as the pdf of the random variable *W*, which is non-negative. Then

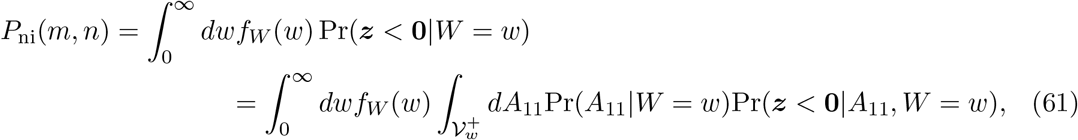

where 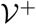 is the set of positive definite symmetric matrices and 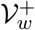 the set conditional to 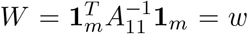. Using that 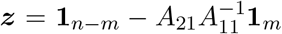 and (60), the conditional variable ***z***|*A*_11_, *W = w* is distributed as

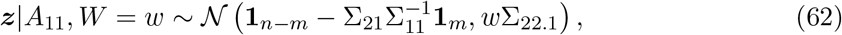

which does not depend explicitly on *A*_11_. Neither does Pr(***z*** < **0**|*A*_11_, *W = w*), so we can factor this probability out of the integration over *A*_11_. In this way, we can write

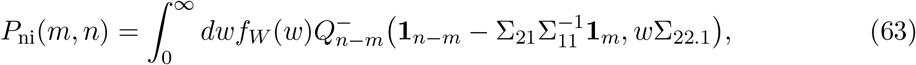

because 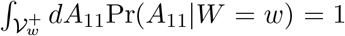. In (63) we have defined 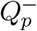 as the probability that a multivariate Gaussian variable with the specified parameters is contained in the fully negative orthant,

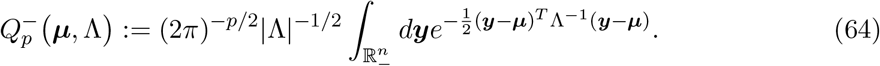

Corollary 3.2.6 in [30] implies that 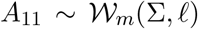. Therefore, theorem 3.2.12 in the same reference holds, which ensures that

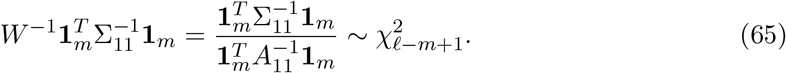

This means that

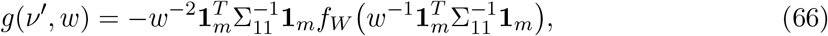

for *g*(*ν, w*) the pdf of a 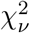, distribution with *ν′ = ℓ − m* + 1 degrees of freedom. Now, making the change of variable 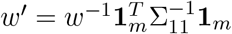 in (63) we finally get

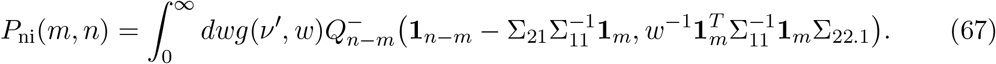

As for the case of feasibility, (67) is an exact formula for the probability that an endpoint composed by *m* species cannot be invaded by the remaining *n − m* species. Similarly, the multidimensional integral associated to 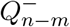 can be reduced to a single integral in the case of constant, non-negative correlation, as we show in the following subsection. Thus, in that particular case, the probability of non-invasibility is expressed as a double integral.

#### Constant, non-negative correlation

In the case of constant, non-egative correlation, (67) simplifies to:

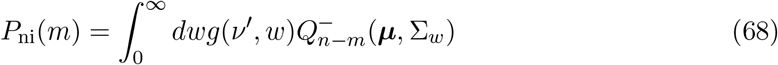

with

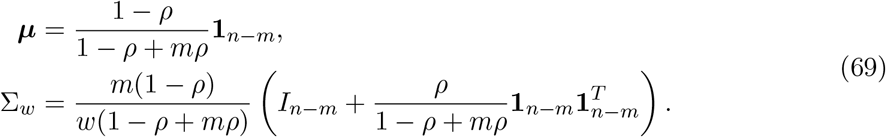

Now focus on the probability 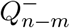. Making the substitution ***y′** = k**y*** in (64) it is easy to show that

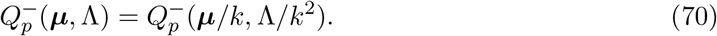

Therefore, for 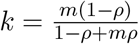 we recover Eq. (84) with ***μ*** and Λ given by

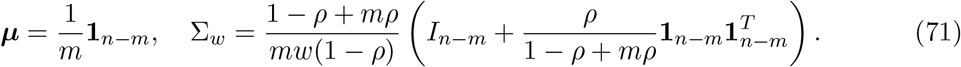

Now let us write 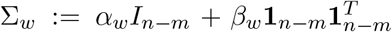, with 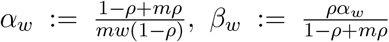. As we did for the probability of feasibility, the probability 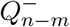 can be written as a one-dimensional integral. For that is crucial that, contrary to what happened in the case of feasibility, correlations given by ∑_*w*_ are positive —notice the plus sign in (71). This is due to the special structure of ∑_*w*_, which implies that the correlation between any two distinct *y_i_, y_j_* in (64) is constant and given by 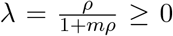. Hence, the following result of Tong [34] (section 8.2.5) applies:

#### Proposition 2.

*Let **x** be distributed according to* 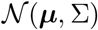 *such that covariance matrix entries satisfy* 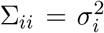 *and* ∑_*ij*_ = *σ_i_σ_j_*λ. *Then, the joint probability that* 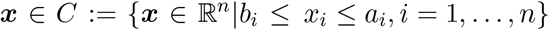, *where* −∞ ≤ *b_i_ < α_i_* ≤ ∞ *for i=1,…, n, is expressed as*

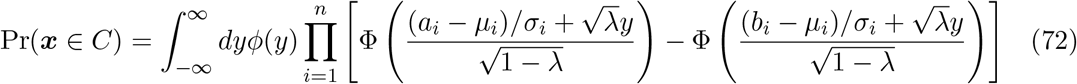

*for φ*(*z*) *and* Φ(*z*) *the pdf and cdf, respectively, of a univariate standard normal distribution.*

In our particular case 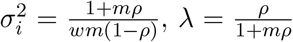, *b_i_* = −∞, *a_i_* = 0 and, according to (71), 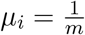 for *i* = 1,…, *n − m*. Therefore, putting all the pieces together, we can write

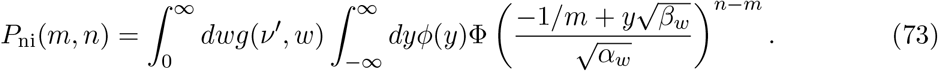

As for the probability of feasibility, in the case of constant, non-negative correlation we can reduce it to a two-dimensional integral.

Notice the resemblance between the expressions for feasibility and non-invasibility — Eqs. (57) and (73). In the case of *ρ* > 0, by changing 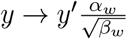, we can make the resemblance stronger:

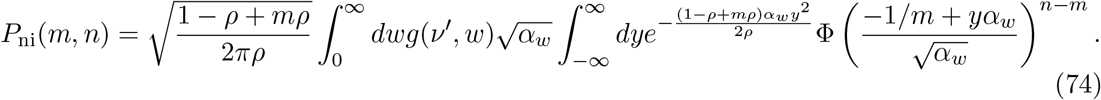

Observe that the number of degrees of freedom of the 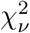, distribution here is *ν′ = ℓ − m* + 1. Notice also that the change of variables leading to (74) does not apply for *ρ* = 0. This case is trivial, however, and will not be discussed explicitly.

#### Sign independence of feasibility and invasibility

In this section we show that the joint probability of feasibility and non-invasibility factors into the product of the two probabilities calculated above. For that purpose, it suffices to show that

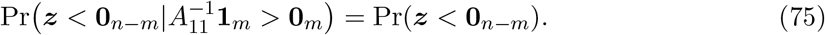

For that purpose we can calculate

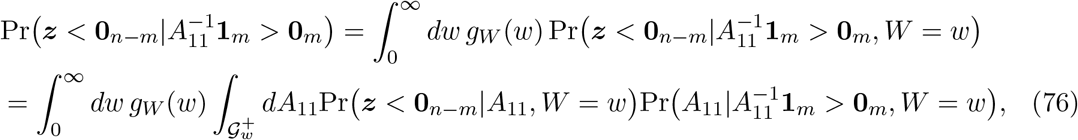

where 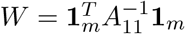 as for the calculation of *P*_ni_, and *g_W_* is the pdf of the random variable 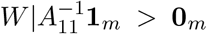. In the second line we have introduced an integral over the set 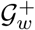 of symmetric matrices and positive definite that verify the conditions 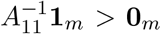 and 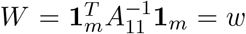. As before, by (62) we can factor the probability Pr(***z*** < **0**_*n−m*_|*A*_11_, *W = w*) out, so we get

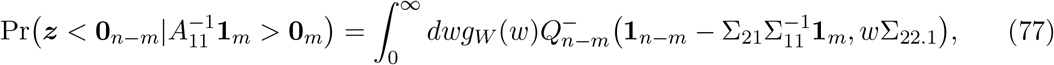

which coincides with (67) except for the probability density *g_W_*. In the last step we have used the normalization condition 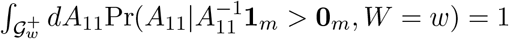.

Observe that the condition 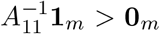 is equivalent to the conditions 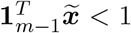 and 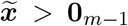, for 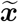 the vector of the first *m* − 1 relative abundances defined in (34). Let 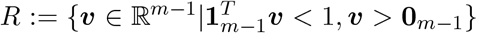 the set of vectors satisfying the two last conditions.

Then it is easy to see that

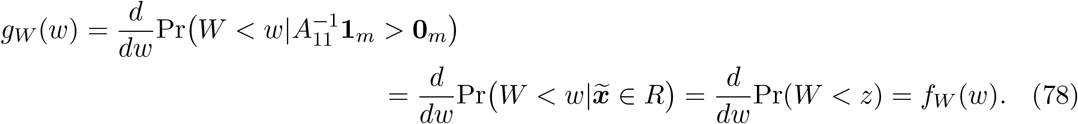

The last equality in the chain above follows because *W* and 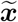 are independent random variables —see the proof of theorem 1 in [8].

This shows that the probability of observing and endpoint with *m* survivors can be factored as the probability of feasibility (37) times the probability (67) that the attractor cannot be invaded by the remaining *n − m* species in the pool.

#### Distribution of the number of coexisting species

Due to the independence shown in the previous section, the probability that the system settles in a subset {*S*}_*m*_ ⊂ {1,…, *n*} formed by *m* species is simply

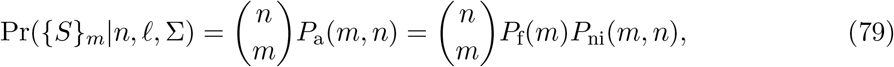

because all subsets with cardinality *m* are statistically equivalent.

Assuming constant and non-negative correlation, in Fig. S5 we compare numerical integration of Eqs. (57) and (73) appearing in (79) with simulations.

#### Average number of species

In this section we will focus on the case of constant correlation. Our aim is to approximate the integrals for feasibility and invasibility in the large number of species limit by a saddle point technique. With these approximations, we provide an analytical way to compute the probability of coexistence Pr({*S*}_*m*_|*n, ℓ, ρ*) —cf. Eq. (79)— as well as an approximation for the average fraction of species

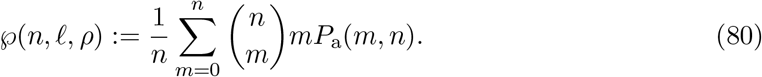

We distinguish the cases *ρ* > 0 and *ρ* = 0 for invasibility. For *ρ* > 0 we use expression (74). Let us define *q* := *m/n* as the fraction of survivors, and recall that *ℓ* = *nγ*. Also let

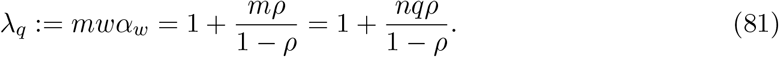

In terms of λ_*q*_, the probability of non-invasibility reads

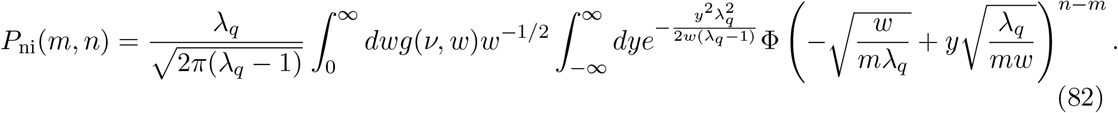

Now we make a change of variables,

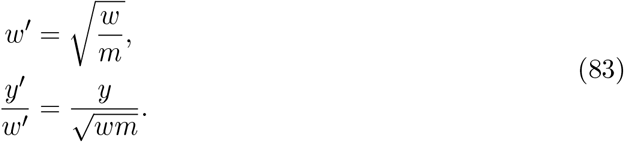

Then the integral becomes

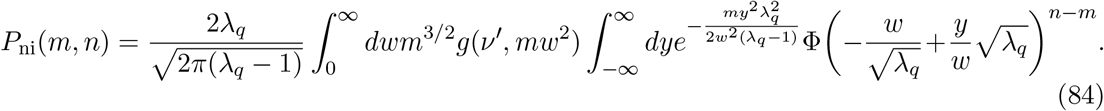

Recall that the probability density function *g*(*ν′,x*), for *ν′ = ℓ − m* + 1, is:

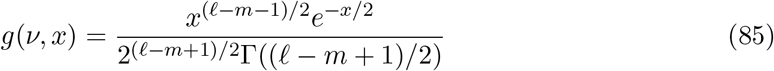

Hence the integral (84) is

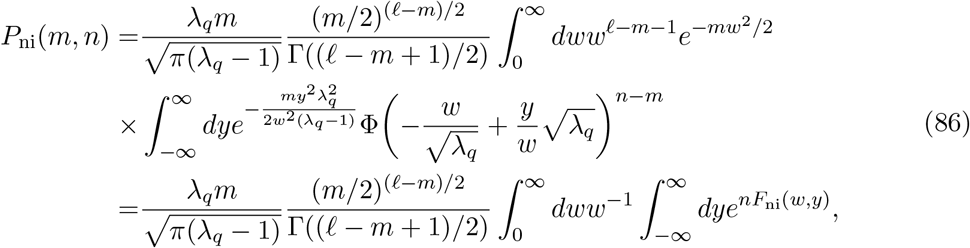

where the exponent *F*_ni_(*w,y*) has been defined as

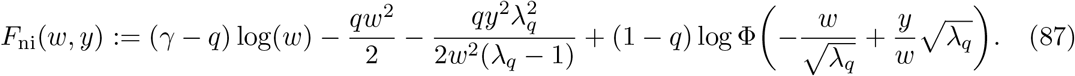

Now we evaluate the double integral in the limit *n* → ∞ via a saddle-point technique. For that purpose, since the exponential becomes peaked around the maximum of the exponent, we calculate the equations to be satisfied by the critical point. Taking derivatives of the exponent we get

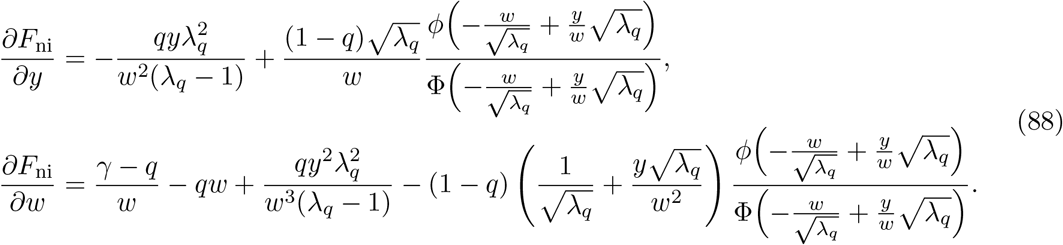

Therefore at a critical point (*w*^★^, *y*^★^) we have the following conditions:

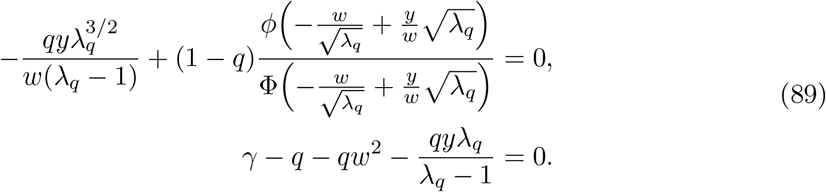

Similarly we can rewrite the integral for the probability that an endpoint formed by *m* species is feasible, see Eq. (57), as

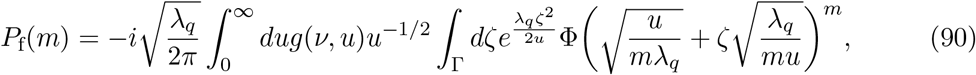

where now the number of degrees of freedom is *ν = ℓ − m* + 2.

Following essentially the same procedure as before, i.e. making a change of variables and replacing the density function for the 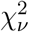 distribution we get

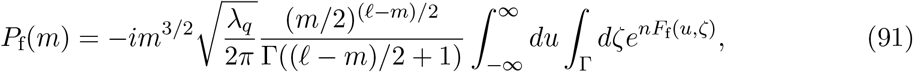

with the exponent

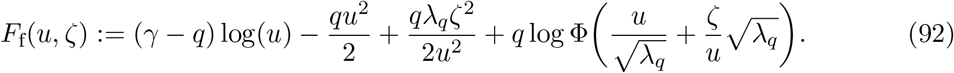

Similarly, the conditions satisfied by the critical point (*u*^★^,*ζ*^★^) are

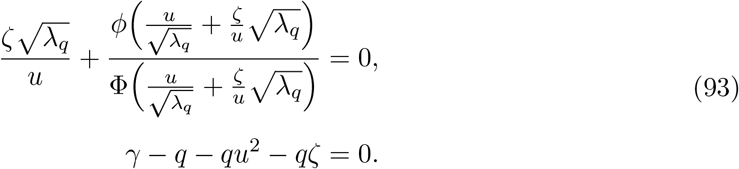

Notice that the product of the densities of the *χ*^2^ distributions in each integral —Eqs. (86) and (91)— introduce an extra term which scales exponentially with *m = nq*, namely

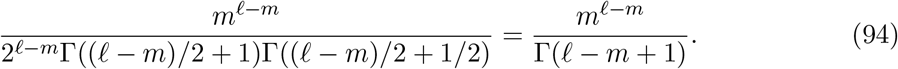

Using the Stirling’s asymptotic form of the gamma function we get

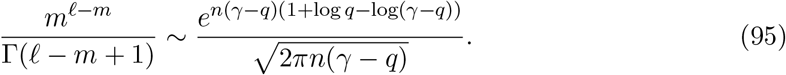

Let

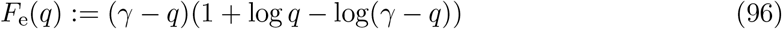

and

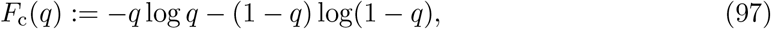

*F*_c_(*q*) being the exponent appearing in Stirling’s asymptotic formula for the binomial coefficient 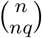. Consequentely the probability that the system settles in an endpoint with *m = nq* species is given, up to a normalization factor, by:

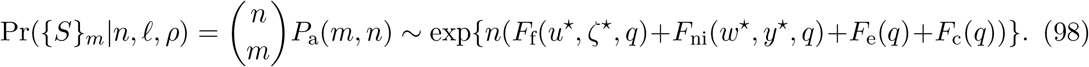

Observe that critical point coordinates *u*^★^, *ζ*^★^, *w*^★^ and *y*^★^ depend implicitly on *q* through (89) and (93). Observe that one can use the asymptotic expansion (98) to obtain numerically the distribution of the number of survivors, Pr({*S*}_*m*_|*n, ℓ, p*), up to a normalization factor. The calculation amounts to solve numerically the non-linear systems (89) and (93).

We are now ready to provide an analytical approximation for the mean fraction of survivors 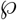, cf. Eq. (80). In the limit of large pool size *n*, we can approximate the mean of the distribution Pr({*S*}_*m*_|*m, ℓ, p*) by its mode, which is easier to compute. In fact, to calculate the mode of the distribution *q* in the large *n* limit we need to find the *q*^★^ value that maximizes the exponent in (98). Due to the critical point conditions for (*u*^★^,*ζ*^★^) and (*w*^★^,*y*^★^), *q*^★^ satisfies

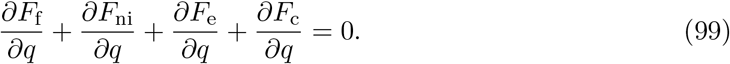

Evaluated at the critical points (*u*^★^,*ζ*^★^) and (*w*^★^,*y*^★^), the derivatives read

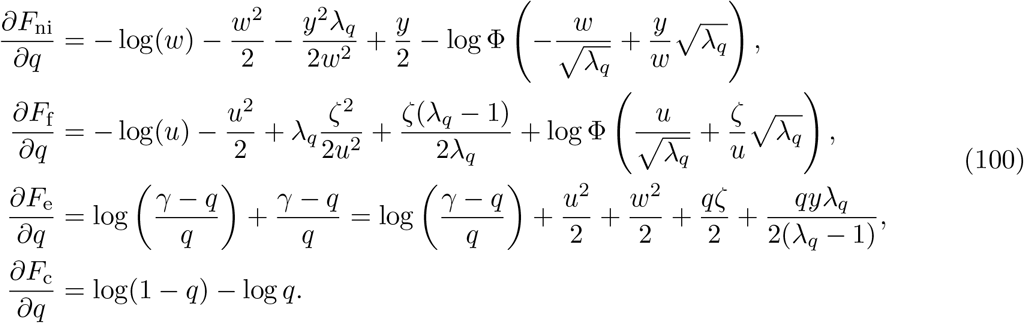

Therefore the condition for *q*^★^ reduces to

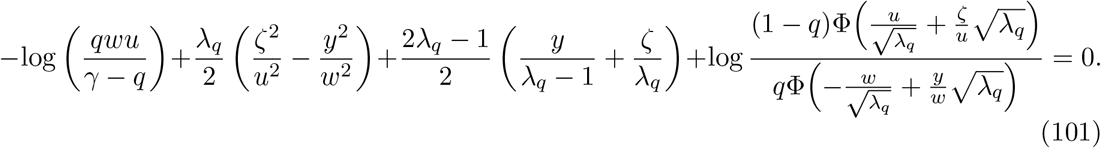

A direct calculation shows that, at 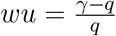, the terms up to the last logarithm vanish. We now show that the last one can be written as 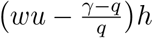 for some function *h*.

Indeed, using conditions (93) and (89) we have

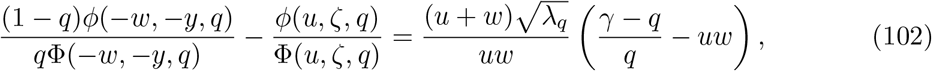

where we have used the abbreviations 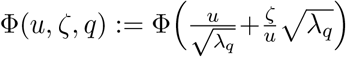 and 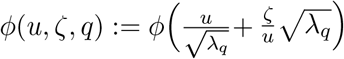 to simplify notation. Therefore,

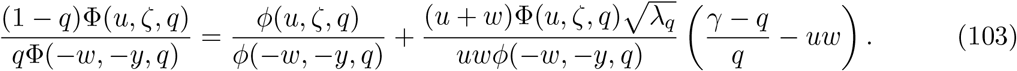

Letting *μ_q_* := (*γ − q*)/*q*, it holds that

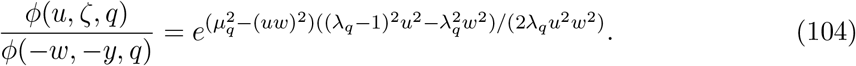

Now, due to the series representation of the exponential function we have

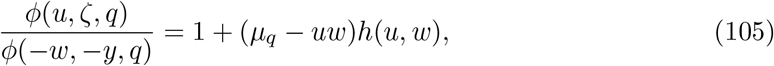

where

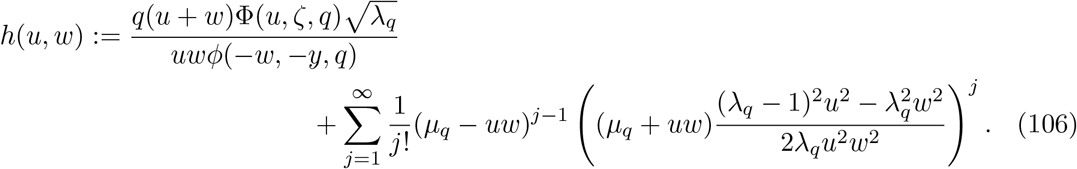

Thus, the claim follows by using the series expansion of log(1 + *x*). Therefore, all the terms in (101) vanish at *uw = μ_q_*.

We have just shown that the last logarithm in (101) is equal to zero. Consequently *q*^★^ satisfies

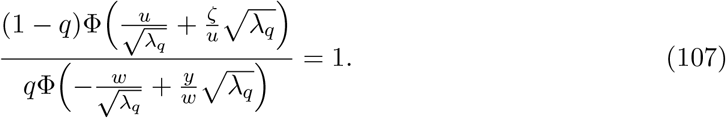

At the point *uw = μ_q_* we can write

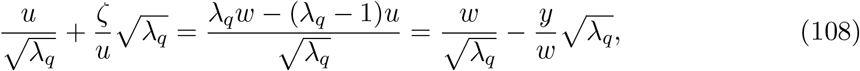

which in turn implies that

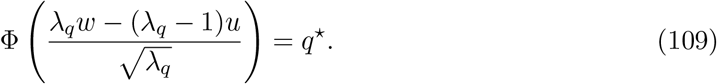

Let 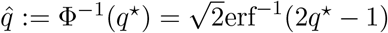, for erf^-1^ the inverse error function. Then it holds that 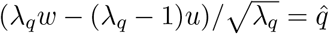 and using eq. (93) we can solve for *u*^★^,*w*^★^ in terms of q, yielding

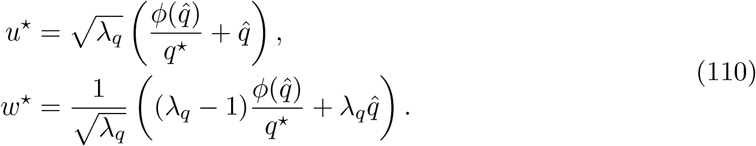

The final condition for *q*^★^ at the saddle point reduces to substitute the expressions above into the condition *uw = μ_q_*, which finally reads

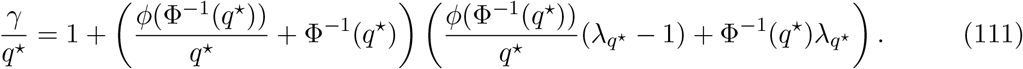

The case *ρ* = 0 for invasibility is similar, and simpler.

#### Level Curves

Eq. (111) gives a very good approximation to the level curves on the (*ρ, γ*) plane mapping to constant mean fraction of survivors *q = m/n*. This implicit condition can be rewritten equivalently as

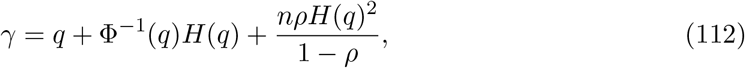

where *H*(*q*) := *φ*(Φ ^−1^(*q*)) + *q*Φ^-1^(*q*). This condition is compared with simulation results in Fig. 3 of the main text (right panel).

### 4 Total biomass distribution at endpoints

The proof of independence of invasibility and feasibility (section 3) also shows that, for any fixed size *m* of a subset of species and total biomass *w*, we have that Pr(***z**_n−m_* < **0**_*n−m*_|***x**_m_* > **0***_m_, W = w*) = Pr(***z**_n−m_* < **0**_*n−m*_|*W = w*). This remark, together with the independence of *W* and ***x**_m_* > **0**_*m*_ (feasibility), helps us derive the distribution of total biomass. To simplify notation we do not rescale the interaction matrix by *ℓ* (as shown in section 6 this would amount to a rescaling of total biomass *w → ℓw*). The cdf for the random variable *W* is precisely

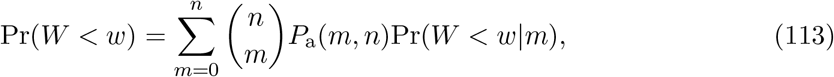

where Pr(*W < w|m*) is the probability that *W < w* conditional on the *m*-species endpoint is feasible and non-invasible. Thus,

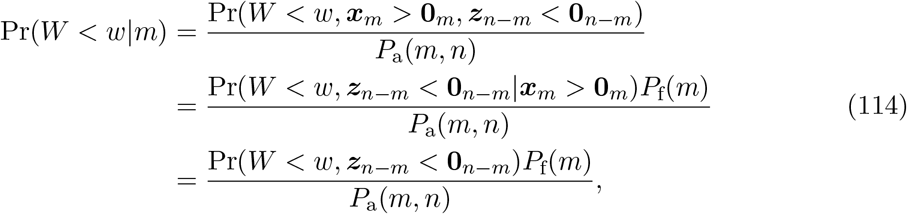

the last equality following from the statement in the paragraph above. Now, using the notations introduced in the last section, it holds that

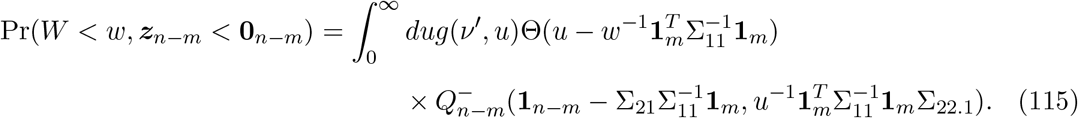

Hence, using (114) and *P_a_*(*m,n*) = *P*_f_(*m*)*P*_ni_(*m, n*), the probability density function of the biomass distribution can be expressed as

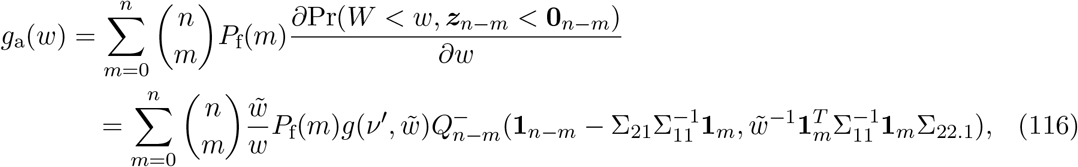

where 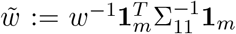. Fig. 12 shows the comparison of (116) with simulations for the constant correlation case in the case in which the interaction matrix is rescaled by the number of traits.

**Figure 12:**
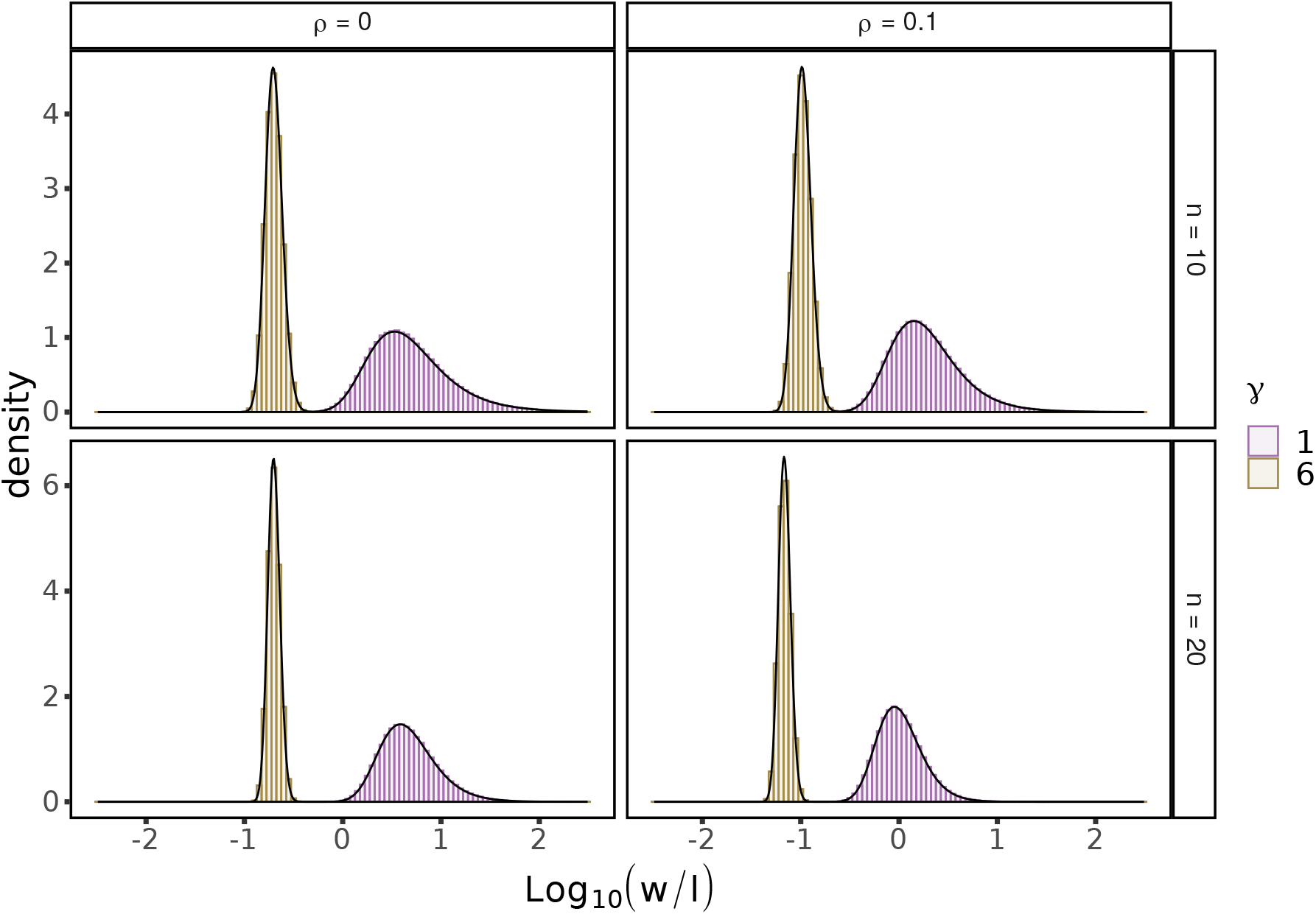
Distribution of the total biomass *w* of the survival community as a function of the ratio *γ* of number of traits *k* to number of species *n* for different *constant* correlation matrices. The simulations were done with *n* = 10, 20 species. Histograms are simulations and black lines are the numerical integration of (116).

**Figure 13:**
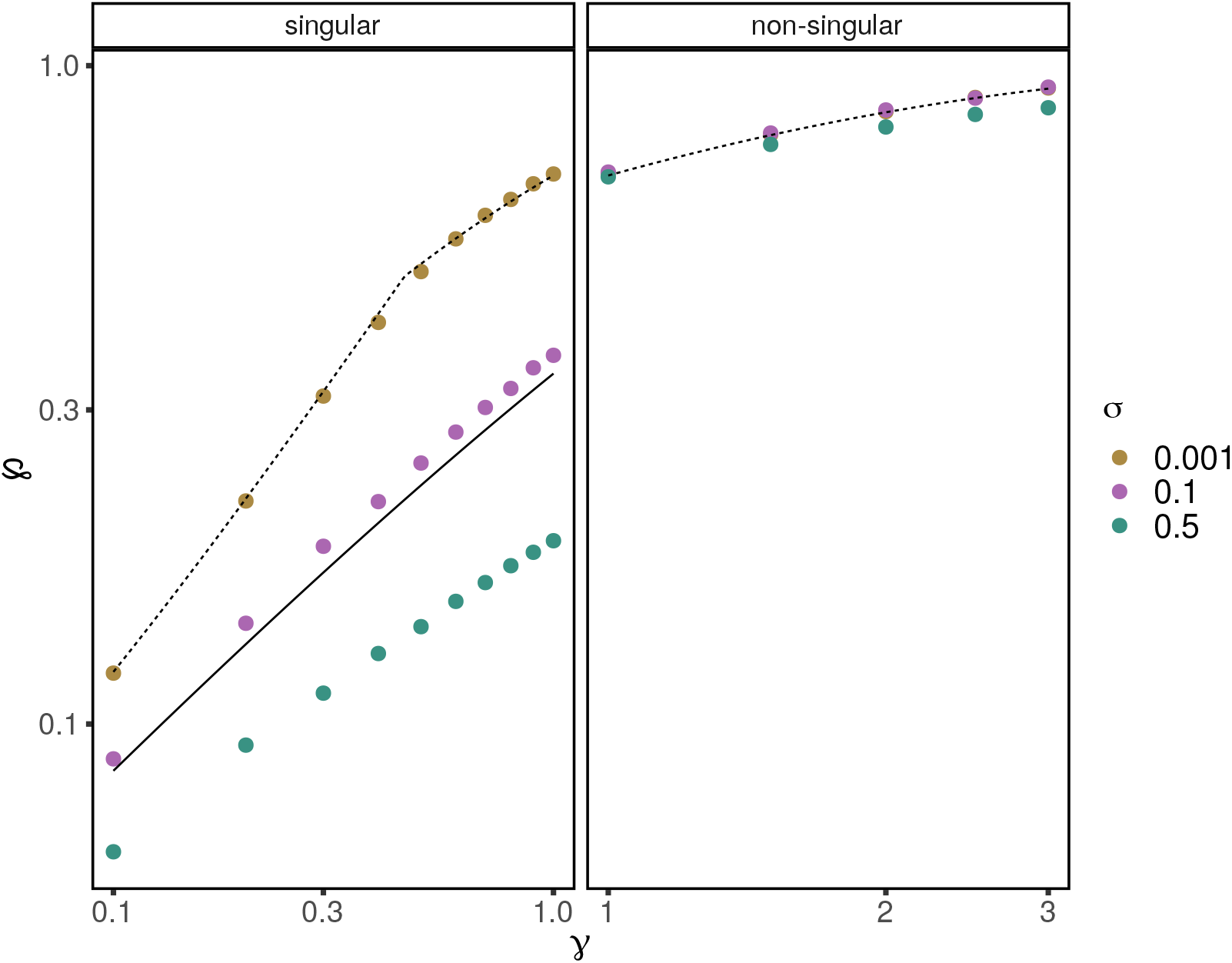
Fraction of survivors under distinct levels of growth rate variability. Dots mark the average values over simulations with 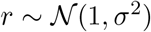 and 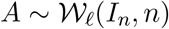. In the singular case, the matrix *A* was perturbed by 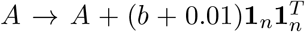 for *b* = −min(*A*). Dotted lines represent our analytical predictions assuming *σ* = 0. By Section 6 the shift in *A* does not affect 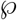 when *σ* = 0. The initial decrease of 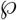 in the singular case is due to this property not holding when *σ* ≠ 0. The solid line is our analytical prediction for *σ* = 0, when 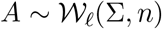. ∑ is a constant correlation matrix with 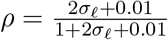 and 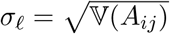 for *i ≠ j* which in this case is simply 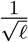.

Going back to re-scaling the interaction matrix by *ℓ*, total biomass transforms as *w → ℓw*. By the above calculation and a change of variables 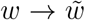, the moments of the distribution of *ℓW* conditional to *m* coexisting species are given by

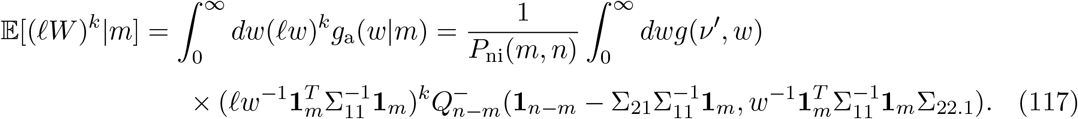

By the saddle point calculation done while computing the expected number of survivors we can approximate the mean of *ℓW|m* for *ρ* ≥ 0, *m* = *nq* and *ℓ = γn* as follows: the above integral satisfies (86) up to a multiplication by 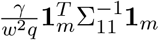 —observe the reescaling in (83). Hence the exponent in the integral does not change so, at the solution (*y*_0_,*w*_0_) of (89), neglecting all but the leading order terms we can approximate

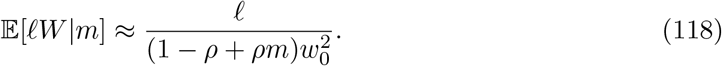

Assuming that the distribution of survivors is highly peaked at the mode, we can approximate the mean of *W* by the mean conditional at the mode, which we get from Eq. (111):

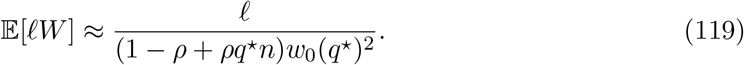

### 5 Relative abundances

For an equilibrium attractor ***x**_m_* with *m* species, let 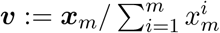 be the relative abundance vector. In particular, 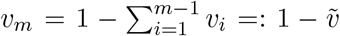. By section 3, Eq. (34), we know that 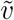 follows a multivariate *t* distribution, so we can write the distribution function for *υ_m_* conditional on ***x*** being feasible as

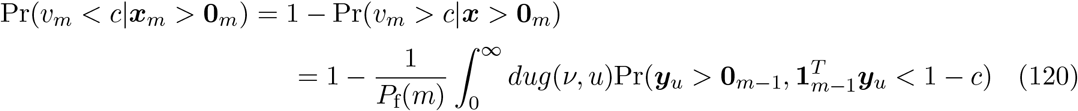

with *ν = ℓ − m* + 2. The independence of 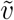 and invasibility gives us the distribution of *υ* conditional to ***x**_m_* being an attractor of the system with *m* out of *n* survivors. Let ***z**_n−m_* be defined as in section 3. Then

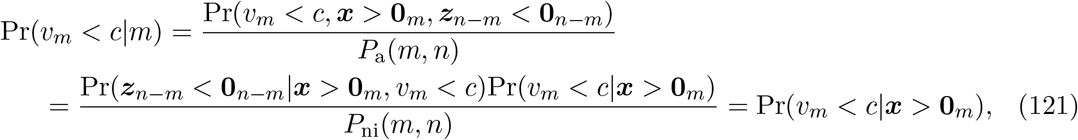

where we have used the independence of feasibility and invasibility, *P_a_*(*m, n*) = *P_f_*(*m*)*P*_ni_(*m, n*).

In case of a constant correlation *ρ* ≥ 0, all species are equivalent so any surviving species *i* has the same distribution as *x_m_*. Applying the same derivation as for the feasibility case, and using the notation of the saddle point calculation with *m = qn* (see Eq. (90)), we get

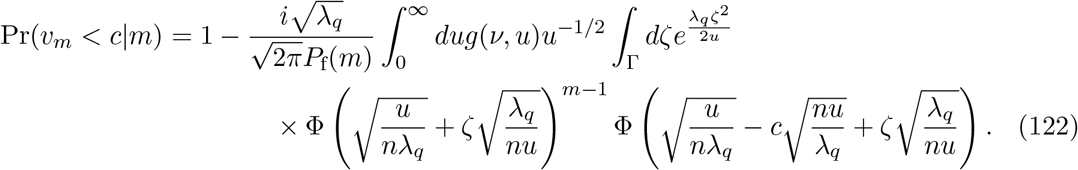

Letting 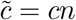, the integral above can be approximated by the same saddle point calculation we did for feasibility (section 3) up to a multiplication factor given by

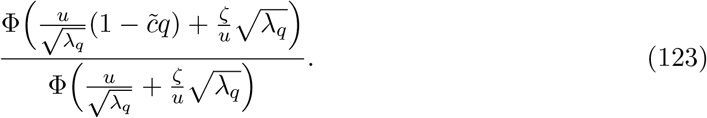

Thus, for (*u, ζ*) satisfying the system of equations (93) with *ζ* real, we get an approximation for the distribution function by neglecting all but the leading terms:

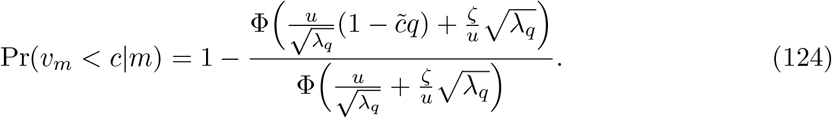

This distribution was compared to simulations in the main text (Fig 4, left panel).

### 6 Invariant Lotka-Volterra operations

In this section we detail the operations that can be performed in a symmetric stable GLV system without changing the subset of coexisting species.

Let 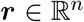 be the vector of growth rates, and 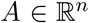 a symmetric and positive definite interaction matrix. Let {*S*}_*m*_ ⊂ {1,…, *n*} be the *unique* subset of *m* species that form the attractor, with vector of densities ***x*** = (*x_i_*). Then ***x*** satisfies:

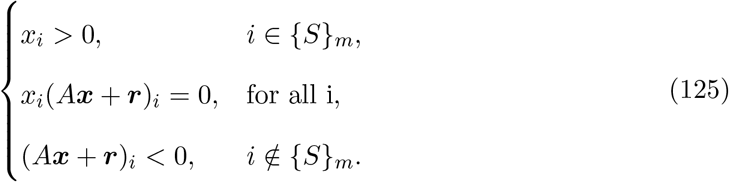

Then we can easily see the effect of the following operations on *A* and ***r*** on the attractor ***x***. Let *κ* > 0 and *D* a positive diagonal matrix. The operations that maintain the identity of the species in the endpoint are:

a. *A → κA*: then ***x** → κ*^-1^***x***.
b. ***r** → κ**r***: Then ***x** → κ**x***.
c. *A → DAD, **r** → D**r***: Then ***x** → D*^-1^***x***.

After any of these operations, the set of coexisting species remains *unchanged*.

Additionally, in the case of ***r** = κ***1**_*n*_, for *κ* > 0, we can perform an additional operation:

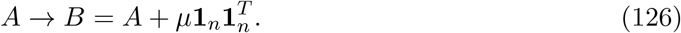

Then shifting

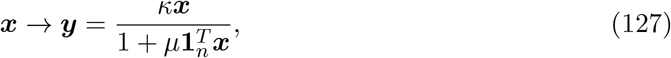

by direct computation of conditions (125) we see that ***y*** is a non-invasible equilibrium. If we additionally restrict *μ* > 0, ***y*** satisfies the feasibility property and *B* is positive definite so again the support {*S*}_*m*_ of the attractor is unchanged.

### 7 Varying growth rates

In this section we analyze the effect that growth rates are not equal for all species. By continuity, we expect our results to hold when ***r*** = **1**_*n*_ + ***ϵ**_n_* and ||***ϵ**_n_*|| ≪ 1 if *ℓ ≥ n*. In case *ℓ < n*, the matrix *A* is singular and the solutions of the system can be unbounded. To correct for that, assume that 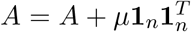 where *μ* is a sufficiently large enough perturbation so that *A_ij_ + μ* > 0. In this case 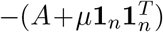 is negative semidefinite and dissipative [19], so the solutions are always bounded. Still, the solutions can be degenerate in the sense that there is a hyperplane of non-invasible equilibria towards which the system converges. By perturbing the growth rates we can correct for that. Assume now that 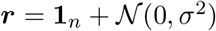, where *σ* ≪ 1 and that 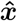 is a saturated rest point of the system (which exists because *A_ij_ + μ* > 0). Without lost of generality, we can assume that the first *m* species survive. Then, we have

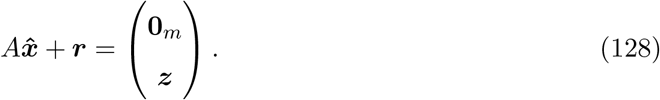

For 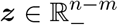, if any *z_i_* = 0, then for the system considering only the species {1,…, *m*} ∪ {*i*} we have that the restriction of ***r*** to this subsystem is contained on a plane of dimension *m* < *m* + 1. Since the distribution of ***r*** is continuous, the probability of this event is 0 almost surely. Hence *z_i_* < 0 for any *i* so that invasibility is *strict*. Furthermore, the same argument shows that the rank of *A* restricted to the survivor subset must be *m*, i.e. the restriction of matrix *A* to the set of coexisting species is *full rank*.

Apply the usual Lyapunov function for the system [19],

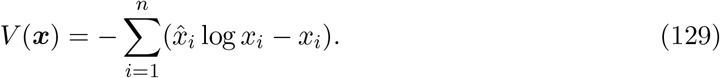

Defined for any 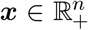, with a global minimum at 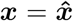 and radially unbounded, then we have

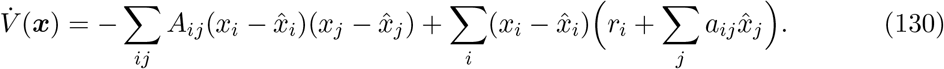

The first sum is non-negative since the matrix is negative semidefinite, and the second is non-positive and is negative unless *x_i_* = 0 for any *i > m*. Given that the restriction of *A* to the survivors subset is full rank then 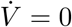 only at 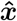, which implies that 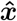 is globally stable and, in particular, is unique [19].

In these cases, while our previous analyses are not exact because of the perturbations introduced in the vector of rates ***r*** and in interaction coefficients 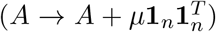, we can apply the same machinery that we have developed to provide approximations. This works because we know that the shift of 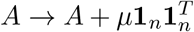 does not change properties like feasibility or invasibility (see section 6). What changes is that the rank of A goes up by one (see the observation below). Forgetting about this, we can use the same machinery as in the non-degenerate case: for feasibility this follows because only full rank subsets are considered, and the restriction of a singular Wishart to a block of *m ≤ ℓ* subsets is a Wishart matrix. Further, the conditional distribution of blocks used for the derivation of the probability of non-invasibility holds in the non-degenerate case too [7].

#### Observation

The rank of 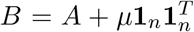 is equal to the rank of *A* plus one. Indeed, let ***w*** ∈ ker *B*, then 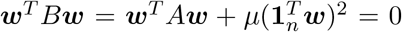, hence ***w*** ∈ ker 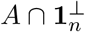, and similarly any ***w*** ∈ ker 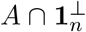 is in the kernel of *B*, hence ker *B* = ker(*A* ⋂ 1^⊥^). Unless ker 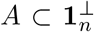, dim(ker *B*) = dim(ker *A*) − 1, so the rank increases by one.

Consider then *A = CC^T^* for 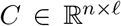, and let {***C**_i_*} be the set of columns of matrix *C*. Then ker *A* is simply *U*^⊥^ = {***C**_i_*}^⊥^. As each column ***C**_i_* is sampled independently from a continuous distribution then *W* = {***C***_1_,…, ***C**_i_*, **1**_*n*_} is a linearly independent set almost surely, then dim *W*^⊥^ = *n* − *ℓ* − 1. Since 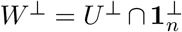, and dim *U*^⊥^ = *n − ℓ* then *U*^⊥^ cannot be contained in 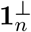.

Observe that the restriction on the size of the subsystems set *γ* + 1/*n* as an upper bound for the mode *q*^∗^. In the singular case it may happen that *q*^∗^ satisfying eq. (111) is bigger than *γ* + 1/*n*. Given that we expect the function to be unimodal and increasing with *q*, then our approximation for the mode in those cases is simply *γ* + 1/*n*.

